# Transgenic mice for *in vivo* epigenome editing with CRISPR-based systems

**DOI:** 10.1101/2021.03.08.434430

**Authors:** Matthew Gemberling, Keith Siklenka, Erica Rodriguez, Katherine R. Tonn-Eisinger, Alejandro Barrera, Fang Liu, Ariel Kantor, Liqing Li, Valentina Cigliola, Mariah F. Hazlett, Courtney Williams, Luke C. Bartelt, Victoria J. Madigan, Josephine Bodle, Heather Daniels, Douglas C. Rouse, Isaac B. Hilton, Aravind Asokan, Maria Ciofani, Kenneth D. Poss, Timothy E. Reddy, Anne E. West, Charles A. Gersbach

## Abstract

The discovery, characterization, and adaptation of the RNA-guided clustered regularly interspersed short palindromic repeat (CRISPR)-Cas9 system has greatly increased the ease with which genome and epigenome editing can be performed. Fusion of chromatin-modifying domains to the nuclease-deactivated form of Cas9 (dCas9) has enabled targeted gene activation or repression in both cultured cells and *in vivo* in animal models. However, delivery of the large dCas9 fusion proteins to target cell types and tissues is an obstacle to widespread adoption of these tools for *in vivo* studies. Here we describe the generation and validation of two conditional transgenic mouse lines for targeted gene regulation, Rosa26:LSL-dCas9-p300 for gene activation and Rosa26:LSL-dCas9-KRAB for gene repression. Using the dCas9^p300^ and dCas9^KRAB^ transgenic mice we demonstrate activation or repression of genes in both the brain and liver *in vivo*, and T cells and fibroblasts *ex vivo*. We show gene regulation and targeted epigenetic modification with gRNAs targeting either transcriptional start sites (TSS) or distal enhancer elements, as well as corresponding changes to downstream phenotypes. These mouse lines are convenient and valuable tools for facile, temporally controlled, and tissue-restricted epigenome editing and manipulation of gene expression *in vivo*.

## Main Text

Epigenome editing with CRISPR-Cas9 systems has become a widespread approach for interrogating fundamental aspects of biological processes through targeted regulation of genes and the non-coding genome.^1^ Fusions of the nuclease-deactivated Cas9 (dCas9) to transcriptional activator or repressor domains enables targeted manipulation of gene expression. Initial work characterizing dCas9 fused to the tetramer of the VP16 acidic activation peptide (dCas9^VP64^) laid the framework for the development of a wide range of next-generation transcriptional activators, including SAM, VPR, p300core, Tet1CD, and others.^2–9^ In addition, dCas9 fusions that can repress gene expression have been characterized to function through direct or indirect chromatin modification using domains and enzymes such as KRAB, MECP2, and DNMT3a.^2, 10, 11^ In particular, we and others have shown that dCas9^p300^ and dCas9^KRAB^ can be targeted to enhancers or near transcriptional start sites, leading to targeted histone acetylation or methylation and subsequent changes in gene expression.^3, 12^ These epigenome modifying dCas9-fusion proteins have been used for studies of gene regulation,^13–15^ directed cell differentiation,^16–20^ therapeutic gene modulation,^21, 22^ and high-throughput screening of putative gene regulatory elements.^23–26^

Although the majority of studies using CRISPR-based epigenome editing tools have focused on *ex vivo* cell culture systems in which delivery challenges are readily addressable, there are several examples of the powerful utility of targeted gene activation and repression *in vivo*. The smaller dCas9 from *Staphylococcus aureus*^27^ was incorporated into a dCas9^KRAB^ fusion protein and delivered with a gRNA expression cassette via adeno-associated virus (AAV) for targeted gene repression in the mouse liver.^21^ Short “dead” gRNAs were used to activate gene expression in the liver and muscle of the Cas9 transgenic mouse^28^ using dual AAV delivery of these gRNAs with an MPH activator module.^29^ A dCas9-SunTag transgenic mouse has been used to activate gene expression in the midbrain^30^ and liver^31^ when a transcriptional activation domain was provided in *trans*. Another study used a constitutively expressed dCas9^VP64^ knock-in to activate expression of the *Sim1* gene and rescue obesity resulting from *Sim1* haploinsufficiency when crossed to mice transgenic for gRNA expression cassettes.^22^ While most previous studies focused on gene activation, one recent study generated a tetracycline-inducible dCas9^KRAB^ mouse line. Hematopoietic stem cells (HSCs) from this line were then engineered *ex vivo* to assess the impact of five transcription factors on hematopoietic lineage determination after transplantation into a host animal.^32^ Plasmids encoding either dCas9^VP64^ or dCas9^KRAB^ have also been transfected into the mouse brain to interrogate the role of epigenetic regulation in addiction behaviors.^13^

These experiments highlight the potential for studies of gene regulation using CRISPR-based epigenome editing. Accordingly, we sought to generate widely applicable transgenic mouse lines to readily perform temporally controlled and tissue-restricted epigenome editing for both gene activation and repression with simple addition of the gRNA. There are several potential advantages to using Cre-inducible expression of dCas9-based epigenome editors from transgenic mouse lines compared to viral delivery. First, several epigenome editors, such as dCas9 fusions to the acetyltransferase p300, are too large for viral vectors. Second, a single genomic copy ensures a more uniform expression level between cells and animals. Third, the use of a genetically encoded Cre recombinase adds tissue or cell type specificity when a corresponding promoter that can be packaged into viral vectors is unavailable. Fourth, it is possible to genetically encode both the gRNA and the Cre recombinase to alter gene expression or program epigenetic states in cell types that are not readily transduced by viral vectors. Fifth, the transgenic expression of dCas9-based epigenome editors may overcome challenges associated with immune system recognition of the bacterial Cas9 protein.^33–37^ Here we characterize and demonstrate the utility of two new mouse lines, Rosa26-LSL-dCas9-p300core (dCas9^p300^) and Rosa26-LSL-dCas9-KRAB (dCas9^KRAB^), for *in vivo* epigenome editing and modulation of gene expression.

### Generation and Characterization of the Mouse Lines

We generated dCas9 epigenome editor mice by inserting a Cre-inducible cassette into the Rosa26 locus using traditional homologous recombination in mouse embryonic stem cells (**Figure 1A**). The inserted transgene consists of a CAG promoter followed by a *loxP*-stop-*loxP* (LSL) cassette, followed by the cDNA encoding the dCas9 fusion protein containing a FLAG epitope. This allows for inducible expression of the dCas9 fusion protein in response to Cre recombinase activity that results in removal of the stop signal between the *loxP* sites. To evaluate the inducibility of the dCas9^p300^ and dCas9^KRAB^ lines, we treated the mice with AAV9:CMV.Cre and found Cre-dependent expression in spleen, skeletal muscle, liver, pancreas, and heart (**Figure S1 and S9B**).

**Figure 1:**
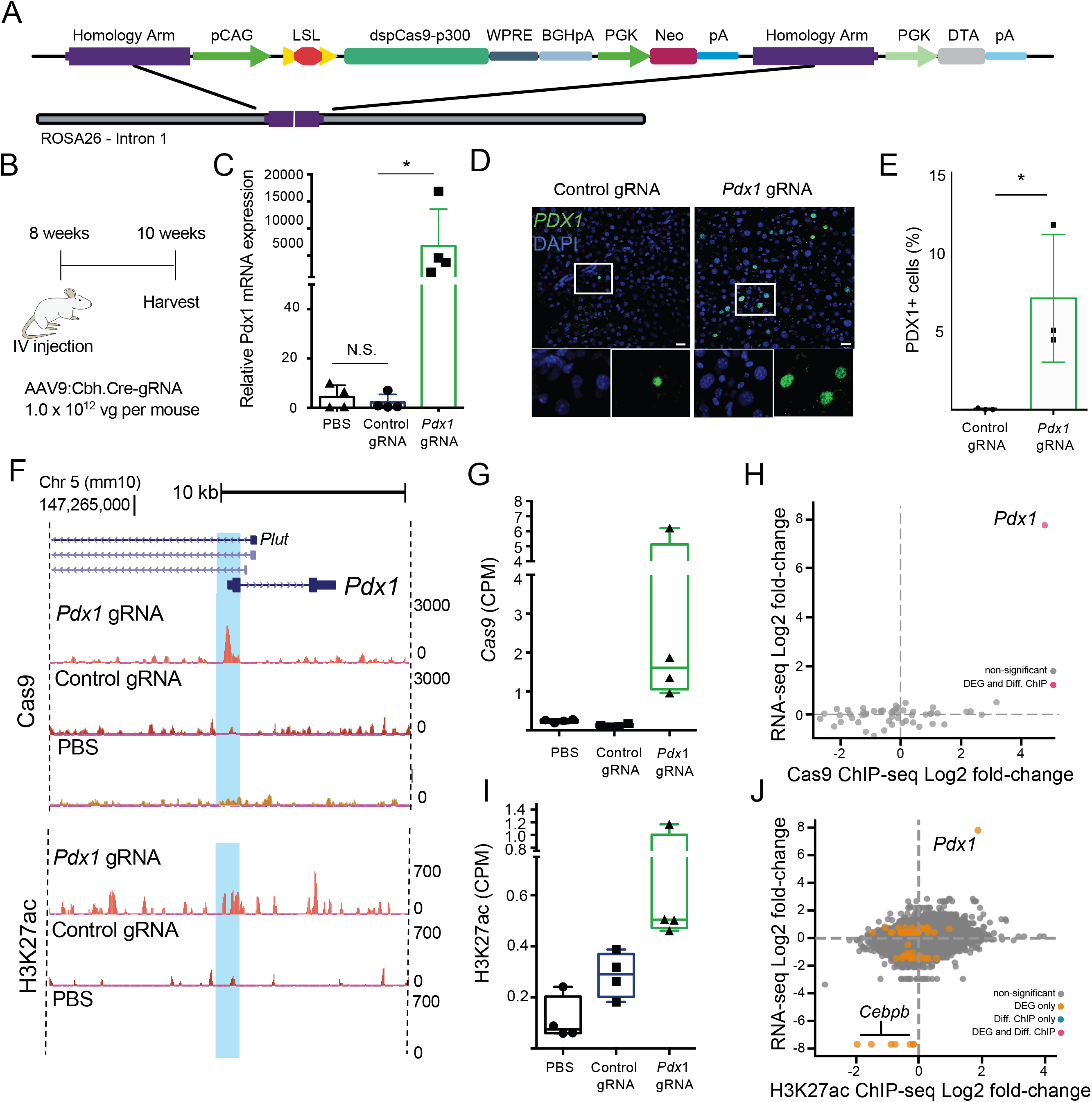
AAV-based gRNA and Cre recombinase delivery to *Rosa26:LSL-dCas9^p300^* mice activates *Pdx1* gene expression and catalyzes targeted histone acetylation. A) Schematic of *Rosa26:LSL-dCas9p300* (dCas9^p300^) knock-in locus. B) Experimental design for *in vivo Pdx1* activation. C) *Pdx1* mRNA quantification 8 weeks post-injection in liver tissue lysates isolated from mice injected with phosphate-buffered saline (PBS), or AAV9 encoding Cre and either a control non-targeting or *Pdx1*-targeting gRNA (n=4 per group, Kruskal-Wallis ANOVA with Dunnett’s post-hoc, *p=0.0132). D) PDX1 immunostaining of liver tissue sections at 14 days post-injection of mice treated with control and *Pdx1*-targeted gRNAs. E) Quantification of PDX1+ nuclei in control gRNA and *Pdx1* gRNA treated animals (7.0% vs 0.03%, p=0.019, students t-test, N=3 animals, with 3 images counted per animal). F) Representative browser tracks of dCas9 and H2K27ac ChIP-Seq data from treated livers at 2 weeks post-treatment (replicates are presented in Figures S3 and S4). G) dCas9 ChIP-seq quantification of sequencing counts (CPM = counts per million) in the gRNA target region of Pdx1 in samples from mice treated with *Pdx1-*targeted gRNA, control gRNA, or PBS (N=4, ANOVA with Dunnett’s post-hoc, p = 0.059). H) Genome-wide RNA-seq and *Pdx1* ChIP-seq analyses shows *Pdx1* gene expression changes observed in *Pdx1* are related to occupancy of Cas9. Log_2_(fold-change) represents the change in read counts between samples treated with the *Pdx1*-targeted gRNA relative to control non-targeting gRNA in dCas9^p300^ cells (N=4, FDR<0.05). I) H3K27ac ChIP-seq quantification of sequencing counts within a 1 kb window centered on the gRNA target site near the TSS of *Pdx1* in samples from mice treated with the *Pdx1*-targeted gRNA, control gRNA, or PBS, show an increase in H3K27ac at the *Pdx1* gRNA target region as compared to controls (N=4, Student’s t-test, p=0.07). J) RNA-seq and H3K27ac ChIP-seq analysis showing significant changes to gene expression are related to increased enrichment of H3K27 acetylation in samples from mice treated with the *Pdx1-*targeted gRNA and control gRNA, although no statistically significant genome-wide differences in H3K27ac levels were observed (N=4, FDR<0.05, DEG = differentially expressed gene, orange dot; Diff. ChIP = differentially enriched ChIP-seq signal, blue dot; DEG and Diff. ChIP = differentially expressed gene and ChIP-seq enrichment, red dot).

### Targeted Gene Activation in the Liver

To demonstrate activation of gene expression with dCas9^p300^ in cells harboring a single genomic insertion of the dCas9^p300^ expression cassette, we isolated primary fibroblasts from the gastrocnemius and tibialis anterior hindlimb muscles of the dCas9^p300^ mouse to test gRNAs in cell culture. We tested four gRNAs targeting the transcriptional start site (TSS) of *Pdx1* (pancreatic and duodenal homeobox 1) (**Figure S2A**). *Pdx1* was originally selected as a target due to its ability to induce liver cells into a pancreatic beta cell-like phenotype, including glucose-responsive insulin expression.^38^ Cells were co-transduced with lentiviral vectors encoding Cre and a single gRNA expression cassette and cultured for 4 days, at which point RNA was isolated for gene expression analysis by qRT-PCR. Two of the four gRNAs significantly increased *Pdx1* mRNA levels compared to a control gRNA targeting *Myod*. For all future experiments, we continued with gRNA #4 as it showed the greatest activation of *Pdx1* mRNA expression at ~150-fold increase relative to controls (**Figure S2B**).

This *Pdx1*-targeting gRNA was cloned into an AAV vector containing a ubiquitous CBh promoter driving Cre (AAV9:Cbh.Cre-Pdx1.gRNA), which we injected into 8 week old mice and collected liver tissue two weeks later to assess *Pdx1* mRNA expression (**Figure 1B**). The *Pdx1* mRNA levels were increased several thousand-fold compared to treatment with a corresponding AAV vector containing a non-targeting control gRNA (**Figure 1C**). We confirmed an increase in PDX1 protein levels in tissue sections of the treated livers as compared to a non-targeting gRNA control as measured by immunofluorescence staining (**Figure 1D**). In the animals treated with *Pdx1* gRNA, 7.0% of nuclei were PDX1+, compared to 0.03% of cells in livers treated with the control gRNA (**Figures 1E**). However, we did not detect an increase in insulin mRNA by qRT-PCR or protein by immunostaining in any treated animals. This result is supported by previous studies showing that hyperactive PDX1-VP16 overexpression was necessary to achieve robust insulin induction.^39^

After demonstrating robust target gene activation, we characterized the specificity of gene regulation in the dCas9^p300^ mice. While the p300 acetyltransferase has many well-characterized targets involved in gene regulation, previous studies showed deposition of acetylation of lysine 27 on histone subunit 3 (H3K27ac) in regions of dCas9^p300^ binding that is concomitant with an increase in target gene mRNA levels.^3, 14, 19, 40–43^ To assess the genome-wide specificity of both dCas9 binding and targeted acetylation, we performed chromatin precipitation and sequencing (ChIP-seq) using antibodies against dCas9 and H3K27ac. Significant levels of specific binding of dCas9^p300^ to the targeted region of *Pdx1* **(Figures 1F-G, S3)** and increased H3K27ac in the surrounding regions were readily detectable (**Figures 1F, S4**). Genome-wide comparisons showed that the only statistically significant differential dCas9 binding site between the *Pdx1*-targeted treatment and non-targeting gRNA controls was at the *Pdx1* gRNA target site (**Figure 1H**). Genome-wide, H3K27ac levels were not significantly different between the treatment and gRNA control **(Figure S5A)**. However, we observed a 2-fold increase in H3K27ac at the *Pdx1* gRNA target binding site as compared to the non-targeting control gRNA **(Figure 1I)**. When comparing mice injected with AAV encoding Cre and the *Pdx1*-targeted gRNA to control mice injected with saline, we found widespread differential H3K27ac levels which appears to be attributable to gRNA-independent consequences of overexpression of the constitutively active p300 acetyltransferase catalytic domain (**Figure S5D**).

To assess the specificity of changes in gene expression downstream of targeted histone acetylation, we performed RNA sequencing (RNA-seq) on mRNA harvested from treated mouse livers. When compared to treatment with Cre and a non-targeting gRNA, the *Pdx1*-targeted gRNA led to increases in *Pdx1* expression that were the greatest change in gene expression transcriptome-wide (**Figure 1H, 1J, S5G, S6A**). This is consistent with previous levels of specificity reported for dCas9^p300^ in cultured cells.^3^ However, when comparing *Pdx1*-targeting gRNA or control non-targeting gRNA to saline-injected controls, we observed widespread gRNA-independent changes in gene expression as a result of dCas9^p300^ expression (**Figure S6B-C**). These results support gRNA-mediated specific epigenome editing and changes to target gene expression *in vivo* in this dCas9^p300^ mouse, but also underscore the need for proper controls to account for gRNA-independent effects when overexpressing constitutively active catalytic domains of epigenome editors.

### Targeted Gene Activation in the Brain

To test gene activation in differentiated neurons with a single genomic insertion of the dCas9^p300^ expression cassette, we first sought to regulate gene expression in cultured primary neurons. Previously, we showed that gRNA-mediated recruitment of dCas9^VP64^ overexpressed from a lentiviral vector is sufficient to drive expression of the mature neuronal NMDA receptor subunit *Grin2c* in developing cerebellar granule neurons (CGNs).^15^ Here, to determine whether the Cre-inducible dCas9^p300^ transgene is similarly effective for activating gene expression, we cultured CGNs heterozygous for the dCas9^p300^ transgene and then either induced dCas9^p300^ expression by lentiviral delivery of *Cre* or over-expressed dCas9^VP64^ from a lentiviral vector for comparison. Recruitment of dCas9^p300^ to the *Grin2c* promoter activated *Grin2c* expression to a similar degree as dCas9^VP64^ recruitment (**Figure S7A**). This activation was specific for *Grin2c* and not due to accelerated neuronal maturation because the expression of another developmentally upregulated gene, *Wnt7a*, was not different in any of the conditions (**Figure S7B**). In addition, these data suggest that the dCas9^p300^ transgene is at least as effective as lentiviral dCas9^VP64^ for inducing expression of developmental genes in cultured primary neurons.

To determine the ability of transgenic dCas9^p300^ to regulate gene expression in neurons *in vivo*, we measured induction of neuronal activity-dependent genes in the hippocampus, a brain region that is important for spatial learning and memory. *Fos* transcription is rapidly and robustly induced by the physiological changes in neuronal firing that follow sensory experience. The *Fos* gene is flanked by five enhancer elements that regulate its stimulus-dependent transcription. We previously showed that expression of dCas9^p300^ from a transfected plasmid and its recruitment to enhancer 2 (Enh2) is sufficient to increase *Fos* mRNA and protein expression levels in cultured neurons.^14^ To recruit dCas9^p300^ to Enh2 *in vivo*, we delivered an AAV vector encoding a gRNA expression cassette, Cre recombinase under the control of the neuron-specific *Syn1* promoter, and GFP to track transduced cells, by stereotactic injection into the dorsal hippocampus of dCas9^p300^ heterozygous mice (**Figure 2A, S8A**). For each mouse, one side of the brain was injected with a gRNA targeting *Fos* Enh2 and the other side was injected with a control non-targeting gRNA against LacZ as an in-animal control. Both sides of the hippocampus showed similar GFP expression and immunofluorescence staining showed robust detection of Cas9 in neurons, confirming that the Cre virus was inducing conditional dCas9^p300^ expression (**Figure S8B**). We allowed one cohort of the mice to explore a set of novel objects in the open field, which is a stimulus that induces expression of *Fos* in the hippocampus.^14^ As expected, immunostaining for FOS protein in the dentate gyrus region of the hippocampus showed sparse cells expressing high levels of induced FOS. However, there was a significantly greater number of high FOS-positive cells on the side of each brain that expressed the *Fos* Enh2-targeting gRNA compared with the side expressing the LacZ control gRNA (**Figure 2B-C**). Importantly, quantification of FOS protein levels across all of the transduced cells showed that the distribution of FOS was significantly increased in the cells expressing the *Fos* Enh2-targeting gRNA compared to those expressing the LacZ-targeting gRNA (**Figure S8C**), consistent with dCas9^p300^-dependent regulation of *Fos* gene expression in a cell-autonomous manner.

**Figure 2:**
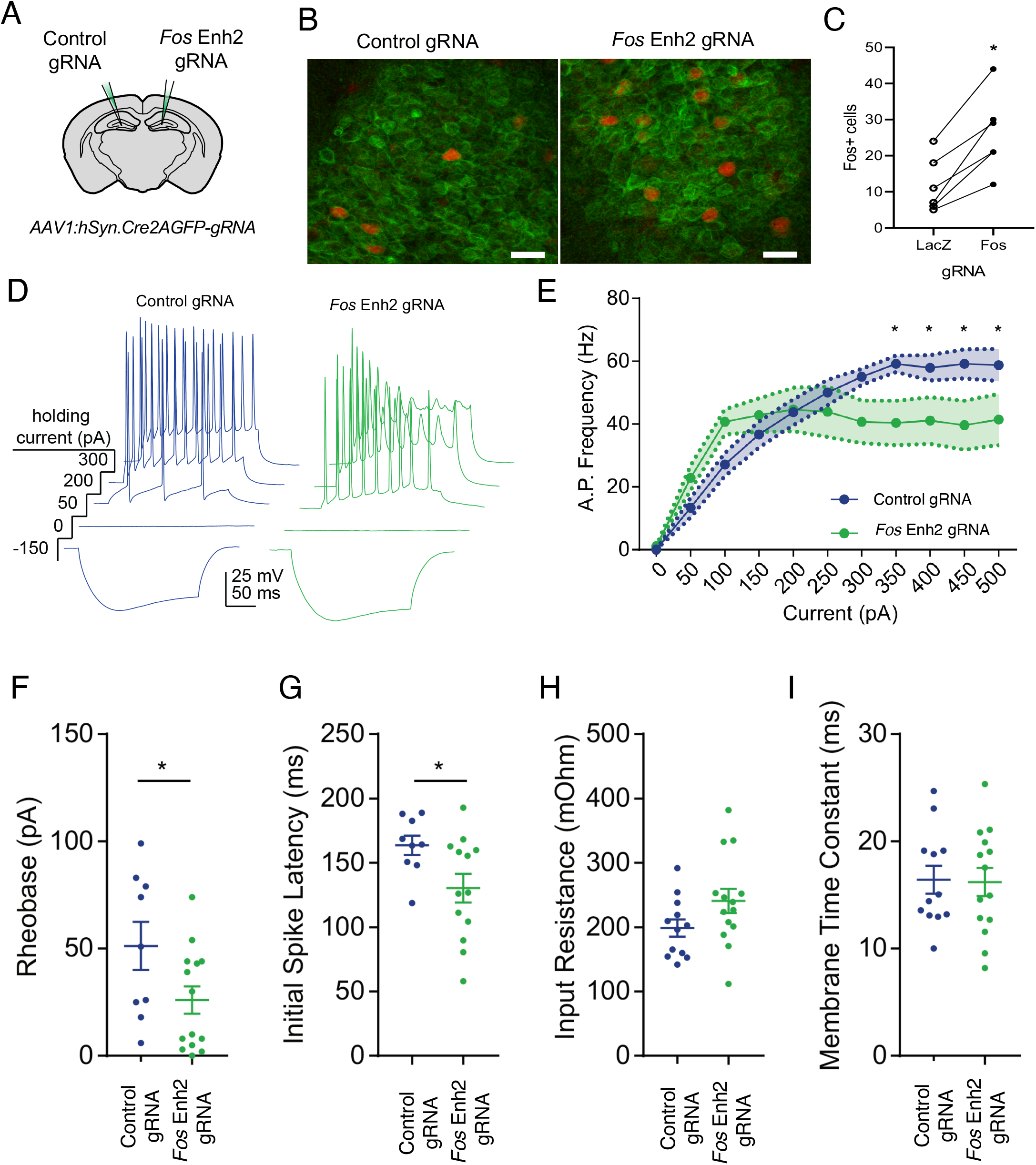
Epigenomic enhancement of *Fos in vivo* increases excitability in CA1 neurons. A) Contralateral AAV injection strategy for comparison of targeting and non-targeting control gRNAs. *AAV:Syn1-Cre.gRNA* was injected into in the hippocampus of dCas9^p300^ mice. A gRNA targeting *LacZ* was used as control for the gRNA targeting *Fos* enh2. B) AAV transduction leads to viral GFP expression in most neurons, and animal exposure to novel objects sparsely induces *Fos* in the hippocampus (scale bar = 20μm). C) Quantification of FOS+ neurons in the dentate gyrus region of the hippocampus after stimulation with novel objects in mice treated with *LacZ* control-gRNA vs. *Fos* Enh2-gRNA. The lines connect measurements from the two sides of the same mouse. Counts of FOS+ cells in tissue slices from n=6 paired ROIs per condition from 3 animals. *p<0.05 by student’s paired t-test. D) Representative current clamp traces from acute hippocampal slices. Virally-expressed GFP was used to identify CA1 neurons for recording. E) Summary of input/output. rmANOVA for current F(10,240)=31.12, p<0.0001, virus F(1,24)=1.05, p=0.32 and current x virus interaction F(10,240)=4.64, p<0.0001. For *Fos* Enh2 vs *LacZ* control at 350pA p=0.016, at 400pA p=0.031, at 450pA p=0.012, and at 500pA p=0.026. F) Rheobase, minimal current to spike. G) Latency to first spike. H) Input resistance. I) Membrane time constant. n=12 *LacZ*, n = 14 *Fos* Enh2, each from 2 animals. *p<0.05 by rmANOVA with posthoc Fisher’s LSD test (E), or paired (C) or unpaired (F-I) student’s t-test. In F-I bars show mean and error bars show SEM.

A key application of using transgenic dCas9^p300^ mice to regulate neuronal gene expression *in vivo* is the ability to determine the consequences of epigenome editing on the function of physiologically relevant neuronal circuits in the intact brain. To determine whether dCas9^p300^-driven increases in FOS protein levels were sufficient to change neuronal physiology, we cut acute slices of hippocampus from a second cohort of dCas9^p300^ mice and performed current clamp recordings from virally transduced CA1 neurons (**Figure S8D-E**). Neurons expressing either the *Fos* Enh2 gRNA or the LacZ gRNA fired action potentials in response to progressive current steps (**Figure 2D**). However, the action potential stimulus-response curves were significantly different between the two treatments. The maximum firing rate was reduced for the neurons expressing the *Fos* Enh2 gRNA compared with neurons expressing the LacZ control gRNA (**Figure 2E**). However, the rheobase, which reports the input current at which neurons begin to fire spikes, and the initial spike latency were both significantly reduced in neurons expressing the *Fos* Enh2 gRNA compared to LacZ gRNA controls (**Figure 2F and G**). These data indicate increased excitability of the *Fos* Enh2 edited neurons. Importantly, the input resistance and membrane time constants between conditions were unchanged (**Figure 2H and I**). This suggests that the differences in action potential firing were due to changes in active, rather than passive, membrane properties. Taken together, these data show that dCas9^p300^-mediated epigenome editing of a single *Fos* enhancer in the adult brain *in vivo* induces changes in FOS protein levels that are sufficient to modulate neuronal physiology.

### Targeted Gene Repression in the Liver

For characterization of the dCas9^KRAB^ mice, we chose to target *Pcsk9* since loss of PCSK9 protein is known to reduce serum levels of LDL cholesterol.^44^ In a previous study, AAV co-delivery of the smaller dCas9 from *S. aureus* fused to the KRAB domain and a *Pcsk9*-targeted gRNA repressed *Pcsk9* in the liver, resulting in reduced serum LDL cholesterol levels.^21^ We delivered AAV9 encoding Cre and an *S. pyogenes* gRNA targeting the same sequence in the *Pcsk9* promoter by tail vein injection to 8-week old dCas9^KRAB^ mice (**Figure 3A**), as well as control mice injected with saline or a corresponding vector containing a non-targeting control gRNA (**Figure 3B**). At eight weeks post-injection, we harvested the mouse livers and assessed *Pcsk9* mRNA levels. We found that *Pcsk9* mRNA levels were reduced ~70% when dCas9^KRAB^ was targeted to the promoter of *Pcsk9* compared to the non-targeting gRNA and saline controls (**Figure 3C**). We also measured serum PCSK9 levels at 4 weeks post-injection and found a ~90% reduction in samples treated with the *Pcsk9*-targeted gRNA as compared to controls (**Figure 3D**). Additionally, serum was collected every two weeks during the eight-week experiment to assess LDL cholesterol levels (**Figure 3B**). There was a ~45% reduction in serum LDL cholesterol levels at each time point from 2 to 8 weeks (**Figure 3E**). Upon *Pcsk9* repression, LDL receptor protein levels increase due to lack of receptor degradation,^45^ which was confirmed by western blot in three of the four Pcsk9-targeted animals (**Figure S9A**).

**Figure 3:**
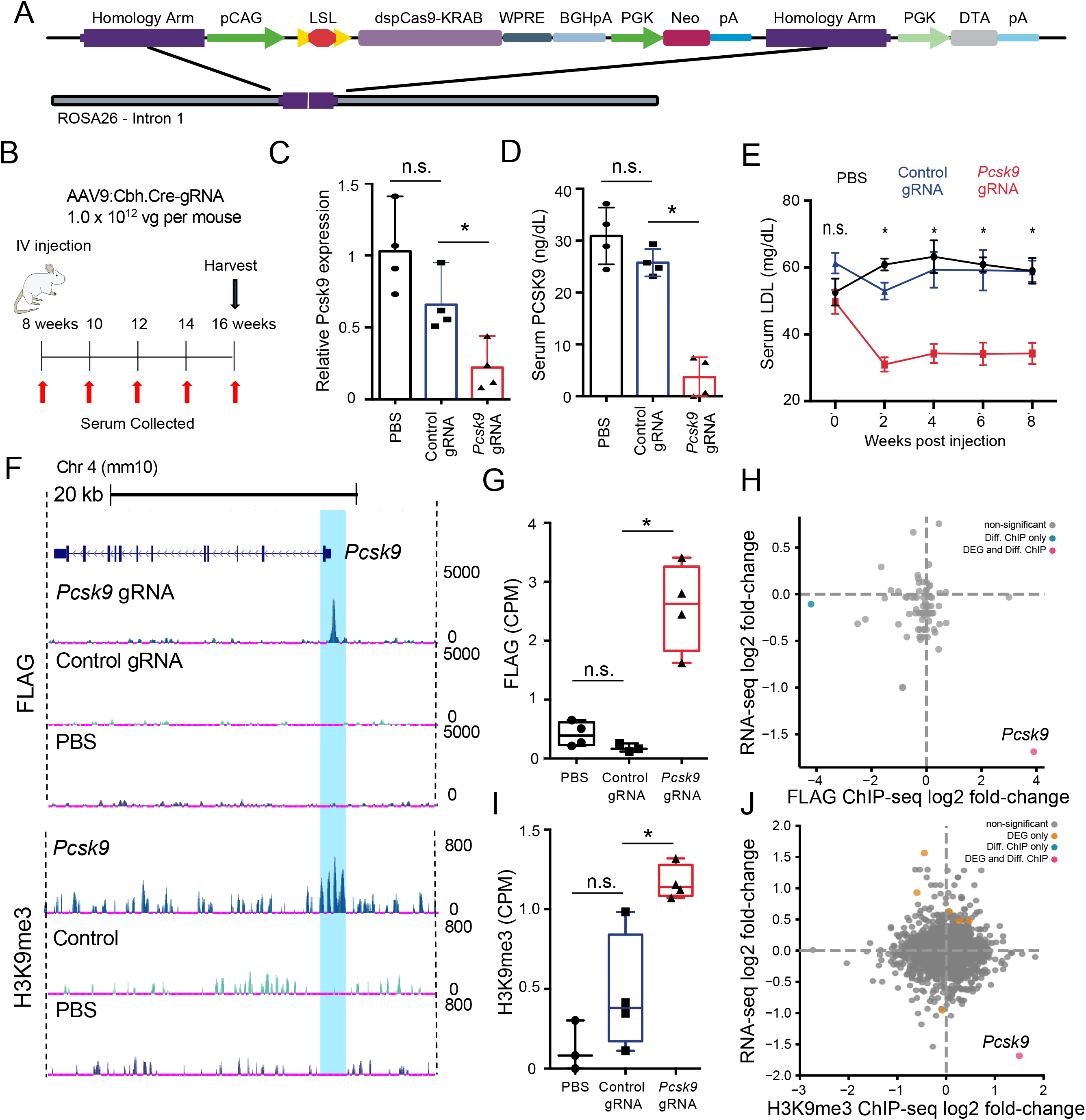
AAV-based gRNA and Cre recombinase delivery to *Rosa26:LSL-dCas9^KRAB^* mice represses *Pcsk9* and catalyzes targeted histone methylation. A) Schematic of *Rosa26:LSL-dCas9KRAB* (dCas9^KRAB^) knock-in locus. B) Schematic of the experimental design to test *Pcsk9* repression in the dCas9^KRAB^ mice. C) *Pcsk9* mRNA quantification 8 weeks post injection in liver tissue lysates isolated from mice injected with phosphate-buffered saline (PBS), or AAV9 encoding *Cre* and either a control non-targeting or *Pcsk9*-targeting gRNA (N=4 per group, ANOVA with Dunnett’s post-hoc, *p=0.0021). D) PCSK9 serum protein levels at 4 weeks post injection of PBS or AAV9 encoding *Cre* and either a control non-targeting or *Pcsk9*-targeting gRNA (N=4 per group, ANOVA with Dunnett’s post-hoc, *p=0.0001). E) LDL cholesterol levels in the serum at 8 weeks following administration of PBS, or *AAV9:Cbh.Cre* containing either a control non-targeting or *Pcsk9*-targeting gRNA (N=4 per group, ANOVA with Tukey’s, *p<0.0001). F) Representative browser tracks of *dCas9^KRAB^* (FLAG epitope) and H3K9me3 ChIP-Seq data from liver samples at 8 weeks post treatment (replicates are presented in Figures S10 and S11). Highlighted region corresponds to the area surrounding the gRNA target site in the *Pcsk9* promoter. G) dCas9 ChIP-seq quantification of sequencing counts (CPM = counts per million) in the *Pcsk9* promoter in samples treated with *Pcsk9*-targeted gRNA, control non-targeting gRNA, or PBS (N=3 or 4 per group, ANOVA with Dunnett’s post-hoc, p < 0.05). H) Log_2_(fold-change) of RNA-seq and FLAG ChIP-seq signal comparing read counts in peaks between samples treated with *Pcsk9*-targeted gRNA and control non-targeting gRNA to highlight the genome-wide specificity of dCas9^KRAB^ binding and function (N=3 or 4 per group, red and blue (*Pcsk9*) dots indicate significant differential peaks, FDR<0.05). I) H3K9me3 ChIP-seq quantification of sequencing counts (CPM = counts per million) in the *Pcsk9* promoter in samples treated with *Pcsk9*-targeted gRNA, control non-targeting gRNA, or PBS (N=3 or 4 per group, ANOVA with Dunnett’s post-hoc, p < 0.05). J) Log_2_(fold-change) of RNA-seq and H3K9me3 ChIP-seq signal comparing read counts in peaks for samples treated with the *Pcsk9*-targeting gRNA and control non-targeting gRNA (N=3 or 4 per group, FDR<0.05, DEG = differentially expressed gene, orange dot; Diff. ChIP = differentially enriched ChIP-seq signal, blue dot; DEG and Diff. ChIP = differentially expressed gene and ChIP-seq enrichment, red dot).

To assess the specificity of gene regulation by dCas9^KRAB^ in these transgenic mice, we performed RNA-seq and ChIP-seq on liver tissue samples from dCas9^KRAB^ heterozygous mice injected with AAV encoding Cre and the *Pcsk9*-targeted gRNA, AAV encoding Cre and the control non-targeting gRNA, or saline controls. For RNA-seq, we performed comparisons of treatment with the *Pcsk9*-targeted gRNA to both the control gRNA and saline controls. In both cases, *Pcsk9* was one of the most strongly downregulated genes, with only 15 and 6 other genes, respectively, significantly changed transcriptome-wide (FDR <0.01, **Figure S6D-F, Supplementary Data**). ChIP-seq was performed using an anti-FLAG antibody for dCas9 binding specificity and a H3K9me3 antibody to detect histone methylation as a result of dCas9^KRAB^-mediated recruitment of methyltransferases (**Figure 3F-J, S10, and S11**). Specific and significantly enriched dCas9 binding to the gRNA target site upstream of *Pcsk9* was readily detected (**Figure 3F-G**). Additionally, significant increases in H3K9me3 deposition in the target region of the *Pcsk9*-targeted gRNA were evident compared to the non-targeting control gRNA or saline controls (**Figure 3F,I**). The dCas9^KRAB^ binding was also highly specific genome-wide, as the *Pcsk9* gRNA target site was one of only two significantly differential peaks between samples treated with the *Pcsk9* gRNA and the control gRNA in the dCas9 ChIP (**Figure 3H**) and the only differential peak in the H3K9me3 ChIP (**Figure 3J**).

### Targeted Gene Activation and Repression in Primary Immune Cells

To further demonstrate the versatility of the dCas9^p300^ and dCas9^KRAB^ mice, we tested modulation of expression of the master regulator transcription factor *Foxp3* in CD4+ T cells. *Foxp3* is essential for the development of a specialized arm of regulatory CD4+ T cells (Tregs) that has a critical role in the attenuation and maintenance of the immune response to self and foreign pathogens.^46–49^ Activation of *Foxp3* expression and the generation of Tregs with immunosuppressive properties is of interest for cellular immunotherapies. Prior work has shown that lentiviral delivery of dCas9^p300^ targeted to the promoter of *Foxp3* was sufficient to increase FOXP3 protein levels.^50^

We activated *Foxp3* in T cells from the dCas9^p300^ mice using a *Foxp3*-eGFP; CD4-Cre; Rosa26-LSL-dCas9^p300^ cross to conditionally express dCas9^p300^ in all CD4+ T cells. Spleen and lymph nodes were isolated from these mice, dissociated, and CD4^+^/CD25^−^/CD44^lo^/CD62L^hi^ naïve T (Tn) cells were purified by fluorescence activated cell sorting (FACS) (**Figure S12A and S12B,C**). Tn cells were cultured in TCR-stimulating conditions (Th0) consisting of plate bound α-CD3, α-CD28, the blocking antibodies α-IL4 and α-IFNg, and IL-2 prior to their transduction with retroviral supernatant containing a gRNA expression cassette. We first tested a panel of 12 gRNAs targeting the *Foxp3* promoter and identified four gRNAs immediately upstream of the TSS which had an activating effect **(Figure S2B, D-E)**. We selected *Foxp3*-gRNA-5 and a non-targeting control gRNA for all downstream experiments. Following retroviral delivery of this *Foxp3*-targeting gRNA, *Foxp3* activation was detected in ~30% of transduced cells as measured by FOXP3-eGFP signal via flow cytometry (**Figure 4A-B**). We verified the gene activation by qRT-PCR and found that the recruitment of dCas9^p300^ to the *Foxp3* promoter led to a ~50-fold increase of *Foxp3* expression relative to the control gRNA or untreated Th0 T cells (**Figure 4C**). Using RNA-seq to assay gene expression in *Foxp3*-gRNA treated Th0 cells, we found that relative to Th0 cells treated with the control gRNA, *Foxp3* was the only differentially expressed gene genome-wide **(Figure 4H, S13)**.

**Figure 4:**
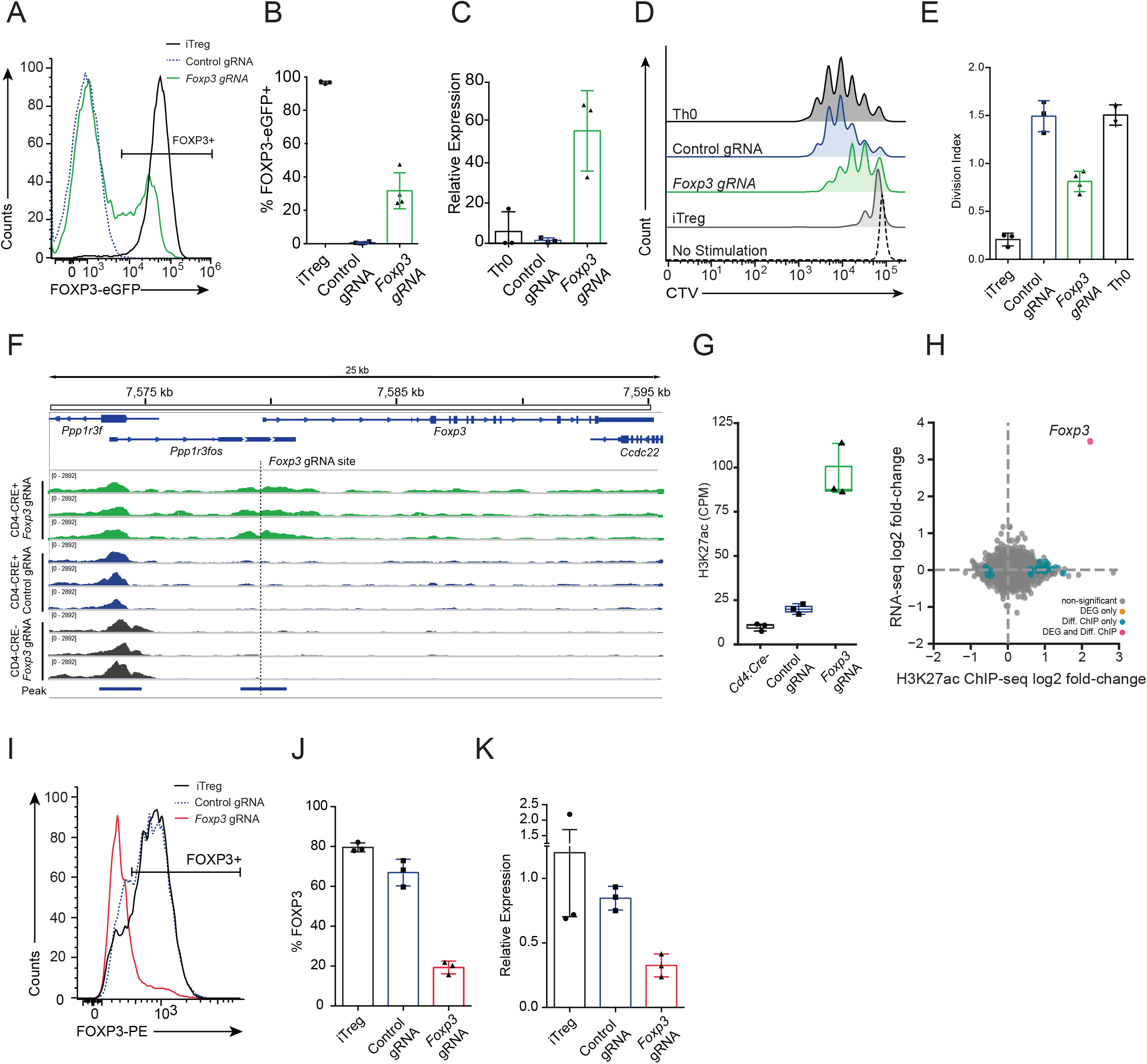
Epigenome editing in T cells for activation and repression of *Foxp3.* A) Flow cytometry analysis of FOXP3-eGFP expression in CD4+ T cells purified from *Rosa26:LSL-dCas9^p300^;CD4:Cre;Foxp3*:*eGFP* mice cultured *in vitro* under Th0 polarization conditions (rIL-2) after transduction with retrovirus encoding the cell surface marker Thy1.1, for tracking transduced cells, and either the *Foxp3-*gRNA (green) or control-gRNA (blue). Cells cultured in iTreg polarization conditions (rIL-2, hTGFβ1) were included as a positive control for *Foxp3* expression (black). B) Summary data depicting the percentage of Thy1.1+ cells that were FOXP3-EGFP+ (p<0.0001. ANOVA with Dunnett’s post-hoc, N=3 or 4). C) qRT-PCR measurement of *Foxp3* mRNA levels in Th0 cells compared to cells treated with *Foxp3*-targeting gRNA (green), control non-targeting gRNA (blue), or no virus (black) (p=0.0190, ANOVA with Dunnett’s post-hoc, N=3). D) Flow cytometry histograms showing proliferation of Cell Trace Violet (CTV)-labelled CD4^+^/FOXP3-eGFP^−^ conventional T cells (Tconv) after 72 hrs of *in vitro* co-culture with aCD3/aCD28 dynabeads and either FOXP3-eGFP+ iTreg cells (grey), Th0 cells (black), FACS-purified Thy1.1+/FOXP3-eGFP+ cells treated with *Foxp3*-gRNA (green), or Thy1.1+ cells treated with control non-targeting gRNA (blue). Tconv cells with no aCD3/aCD28 served as a no activation control (dashed). E) Suppressive capacity of Tregs summarized as division index of Tconv (p<0.0001 ANOVA with Tukey’s post-hoc, N=3 or 4; Th0, N=2). F) Browser track of H3K27ac ChIP-seq read counts at the *Foxp3* locus in Th0 cells from a *Rosa26:LSL-dCas9^p300^* mouse that was either *Cd4:Cre*+ or *Cd4:Cre*−. Cells were treated with retrovirus containing *Foxp3-*targeting or control non-targeting gRNA and *Thy1.1* reporter. ChIP-seq was performed 72hrs after transduction on sort-purified Thy1.1+ cells (red = *Cd4:Cre+ Foxp3*-gRNA; yellow = *Cd4:Cre+* control-gRNA; blue = *Cd4:Cre-Foxp3*-gRNA). G) Summary of H3K27ac ChIP-seq reads counts per million within the MACS2-called peak that intersects the *Foxp3*-gRNA target site for each genotype and gRNA treatment (green = *Foxp3*-gRNA treated *Cd4:Cre*+ Th0, blue = control-gRNA treated *Cd4:Cre*+ Th0; black = *Foxp3*-gRNA treated *Cd4:Cre*-Th0) H) Scatter plot depicting log_2_(fold-change) of gene expression and H3K27ac enrichment when comparing read counts from *Cd4:Cre+ Rosa26:LSL-dCas9^p300^* Th0 cells treated with *Foxp3*-gRNA to control-gRNA (FDR < 0.01; DEG = differentially expressed gene, orange dot; Diff. ChIP = differentially enriched ChIP-seq signal, blue dot; DEG and Diff. ChIP = differentially expressed gene and ChIP-seq enrichment, red dot) I) Flow cytometry analysis of FOXP3 expression in CD4+ T cells purified from *Rosa26:LSL-dCas9^KRAB^* mice cultured *in vitro* under iTreg polarization conditions and transduced with retrovirus encoding the indicated gRNAs. J) Summary data showing the percentage of Thy1.1+ cells that were FOXP3-EGFP+ for each gRNA treatment (p<0.0001, ANOVA with Dunnett’s post-hoc, N=3). K) qRT-PCR measurement of *Foxp3* mRNA levels in iTreg cells treated without virus (black), control-gRNA (blue), or *Foxp3*-targeting gRNA (red) (p<0.0135, ANOVA with Dunnett’s post-hoc, N=3).

To quantify changes in genome-wide histone acetylation made by dCas9^p300^, we performed ChIP-seq for H3K27ac in Th0 cells from *Cd4-Cre+* dCas9^p300^+ mice transduced with *Foxp3*-targeting or control gRNAs. We identified a significant enrichment in H3K27ac at the gRNA target site and only 25 other H3K27ac peak regions were detected as differentially acetylated genome-wide **(Figure 4F-H**, **Figure S5B**, **Supplementary Data**). Together, the RNA-seq and ChIP-seq analyses highlight the specificity and efficacy of targeting dCas9^p300^ to one location of the genome (**Figure 4H**). However, when we compared RNA-seq and ChIP-seq in Th0 cells treated with the *Foxp3*-gRNA from mice with or without *Cd4-Cre*, we observed numerous differences in gene expression and H3K27 acetylation **(Figure S5H, Supplementary Data)**. When comparing RNA-seq data from Cre- vs Cre+ cells, the same set of genes were found to be differentially expressed regardless of whether cells were treated with *Foxp3*-targeting or control gRNA, further supporting a guide-independent effect of active dCas9^p300^ expression on the baseline epigenetic state in this mouse strain **(Figure S13)**. This effect is specific to the dCas9^p300^ mice and is not observed in dCas9^KRAB^ mice when comparing liver samples from mice treated with either *Pcsk9*-targeting gRNA or saline **(Figure S5F,I)**. These results reiterate the need for experimental controls that include an active dCas9^p300^ for comparisons to an accurate baseline epigenetic state.

To test the capacity of FOXP3+ Th0 cells generated by targeted epigenome editing to function as Tregs, we used an *in vitro* suppression assay to test their ability to limit proliferation of activated CD4^+^ T cells in co-culture. The cells treated with *Foxp3*-targeting gRNA showed enhanced suppression relative to untreated cells or cells treated with control gRNA controls, showing suppressive activity similar to the positive control iTregs generated *in vitro* by established protocols **(Figure 4D and E)**. To verify that the suppression activity was mediated by the gRNA-treated cells, we performed the suppression assay with serial dilutions of the number of *Foxp3-*induced cells in co-culture and found a dose-dependent suppressive effect (**Figure S12F and G**).

To demonstrate gene repression in another cell type from the dCas9^KRAB^ mice *ex vivo*, we isolated naïve CD4+ T cells from the lymph nodes and spleen of Rosa26-LSL-dCas9^KRAB^ Cd4:Cre mice and generated FOXP3+ iTregs *in vitro* via TCR activation and treatment with IL-2 and TGFβ1. iTregs were transduced with retrovirus containing either the *Foxp3-*targeting or non-targeting gRNAs. We achieved ~70% reduction in *Foxp3* mRNA levels in cells with dCas9^KRAB^ targeted to the *Foxp3* promoter as compared to untreated iTregs or those treated with the control gRNA (**Figure 4I**). In addition, we assessed FOXP3 protein levels by immunostaining and found a significant reduction in the percentage of FOXP3+ cells when transduced with the *Foxp3*-targeting gRNA as compared to controls (**Figure 4J-K**). These results confirm our ability to repress genes in multiple tissues and cell types of the dCas9^KRAB^ mouse line both *in vivo* and *ex vivo*.

## Discussion

Perturbation of gene expression has long been a powerful tool for uncovering the functions of genes. Recent technological advances have enabled not only the study of gene function, but also the mechanism by which the genes themselves are regulated. Using dCas9 fused to epigenome modifiers has proven to be a productive strategy to uncover unknown gene function, to map regions of the non-coding genome, and to validate potential therapeutic applications using CRISPR-based epigenome editors *in vivo*. Here we describe the generation and characterization of Cre-inducible dCas9^p300^ and dCas9^KRAB^ transgenic mouse lines for targeted activation or repression of promoters and non-coding regulatory elements *in vivo* or in primary cells *ex vivo*, including in the liver, T cells, fibroblasts, and neurons. We demonstrate induction of dCas9 epigenome editor expression using viral or transgenic Cre delivery combined with viral gRNA delivery. The targeted gene activation or repression induced both changes in mRNA transcript levels and protein changes that elicit downstream phenotypes. Concomitant with expression changes, we show targeted deposition of histone marks subsequent to the recruitment of dCas9-based epigenetic effectors at both promoters and enhancers.

An unbiased, comprehensive, genome-wide analyses of epigenome editing specificity by ChIP-seq and RNA-seq showed both a high level of precision in DNA-targeting and interesting differences between the dCas9^p300^ and dCas9^KRAB^ editors. The dCas9^p300^ editor was highly precise and robust in binding to its genomic target (**Figure 1H**), which led to corresponding specificity in catalyzing H3K27ac (**Figure 1F-J, 4F-H**) and activating target gene expression (**Figure S5A, B**) when compared to a control non-targeting gRNA. However, when compared to saline injection or *Cd4-Cre*- control samples, in which Cre is not present and the dCas9^p300^ would not be induced, there were widespread genome-wide changes to H3K27ac and gene expression (**Figure S5G, H**). Interestingly, these changes did not appear to result in any adverse phenotype. Since the off-target changes were similar between the targeting and non-targeting gRNA in T cells (**Figure S13**), we interpret the changes to be gRNA-independent and related to the expression of the constitutively active core catalytic domain of the p300 acetyltransferase that may alter the genome-wide epigenetic baseline of these cells. This is consistent with previous observations that dCas9 fused to catalytic enzymes such as DNA methyltransferases can exhibit gRNA-independent changes,^51^ and underscores the importance of using proper gRNA controls to determine the consequences of modulating gene expression through epigenome editing. In contrast to dCas9^p300^, which has inherent non-specific catalytic activity, dCas9^KRAB^ serves as a scaffold to recruit other enzymatic histone modifiers. Accordingly, target changes to gene expression and epigenetic state were highly specific when comparing to either the non-targeting gRNA or saline controls (**Figure S5C, F, I**), which was consistent with the high level of specificity of DNA-targeting (**Figure 3H**) and histone modifications (**Figure 3J**).

Following the development of CRISPR-Cas9 for genome editing in human cells,^52–56^ a Cre-inducible Cas9 transgenic mouse was quickly generated and widely distributed for *in vivo* studies of gene function genetics of disease.^28^ This mouse line has been essential to numerous important breakthroughs including *in vivo* dissection of gene function^57^ studies of gene regulatory networks,^58^ and *in vivo* genetic screens.^59^ We expect these mice to be similarly powerful for enabling *in vivo* studies of endogenous gene knockdown, activation, and perturbation of epigenetic states. Accordingly, the mice have been deposited for distribution via The Jackson Laboratory (Stock No: 033065 (dCas9^p300^) and 033066 (dCas9^KRAB^)).

## Acknowledgements

We thank Jay Deng and Brendan Ryu for assistance with viral injections and Kevin Franks for his support in performing the electrophysiology recordings. This work has been supported by the Translating Duke Health Initiative, the Allen Distinguished Investigator Program through The Paul G. Allen Frontiers Group, Open Philanthropy, National Institutes of Health (NIH) grants R33DA041878, R01DA036865, U01AI146356, UM1HG009428, UG3AR075336, and R01GM115474, National Science Foundation (NSF) grant EFMA-1830957, and Defense Advanced Research Projects Agency (DARPA) grant HR0011-19-2-0008. V.C. was supported by a Swiss National Science Foundation Postdoctoral Fellowship.

## Author Contribution

M.G., K.S., E.R., K.T-E., F.L., A.K., V.C., M.F., L.B., C.W., and J.B. conducted experiments and analyzed data. H.D., C.R. and L.L assisted with mouse experiments. A.B., and K.S. performed ChIP-Seq and RNA-seq analysis. M.G., K.S., A.W., and C.G. wrote portions of the manuscript. I.H. provided critical reagents. V.J.M, and A.A. produced AAV9 for mouse experiments. M.C., K.P., T.R., A.W., and C.G. provided guidance for experimental design and interpretation of results. All authors edited the text.

## Materials and Methods

### Generation of Rosa26:LSL-dCas9-p300 and Rosa26:LSL-dCas9-KRAB transgenic lines

All experiments involving animals were conducted with strict adherence to the guidelines for the care and use of laboratory animals of the National Institute of Health (NIH). All experiments were approved by the Institutional Animal Care and Use Committee (IACUC) at Duke University. To generate mouse lines for conditional expression of these dCas9 fusion proteins, we used a modified pAi9 targeting vector. The pAi9 vector targets the Rosa26 locus and contains the following features: 5’ Rosa homology arm, CAG promoter, a loxP flanked triple polyadenylation signal (pA) stop cassette (lox-stop-lox; LSL), a codon optimized dCas9-p300 cassette (Hilton et al., 2015) or dCas9-KRAB (Thakore et al., 2015) with either a 1X-FLAG or 3X-FLAG respectively, a woodchuck hepatitis posttranscriptional regulatory element (WPRE), a bovine growth hormone pA, a PGK-Neo-pA selection cassette, and the 3’ homology arm. This modified pAi9 targeting vector was electroporated into hybrid G4 B6N/129S6 ES cells and targeting of the ROSA locus was confirmed by PCR and sequencing. Positive clones were expanded and injected into the 8-cell morulae of ICR mice. Chimeric mice were then mated to establish the transgenic line. Mice are genotyped using the following primers: Forward Primer: GCAGCCTCTGTTCCACATACAC, Reverse Primer: TAAGCCTGCCCAGAAGACTC, Second F primer: AAAGTCGCTCTGAGTTGTTAT. The following PCR conditions are used for genotyping PCR: 95°C for 5 min, 35 cycles of: 95°C for 30 seconds, 57°C for 45 seconds, 72°C for 1 minutes, 72°C for 3 minutes, and 12°C Hold. Expected product sizes are as follows: WT band is 235bp; Knock-In is 162bp. All Mice used in experiments were dCas9^p300^ or dCas9^KRAB^ heterozygous animals.

The new mouse lines are available via The Jackson Laboratory: Stock No: 033065 | Rosa26-LSL-dCas9-p300; Stock No: 033066 | Rosa26-LSL-dCas9-KRAB.

### Isolation and culture of primary dCas9^p300^ cells

Fibroblasts were isolated from gastrocnemius and tibialis anterior of hindlimbs of 6 week old dCas9^p300^ heterozygous animals using a modified version of a protocol by Springer et al (Springer et al., 2002). Instead of proceeding with myoblast isolation, fibroblasts were selected through selective trypsinization of the myoblasts out of the culture using 0.25% Trypsin. Fibroblasts were then grown in DMEM +10% FBS.

Cerebellar granule neurons (CGNs) from male and female postnatal day 7 (P7) dCas9^p300^ heterozygous pups were cultured following our published protocols (Frank et al., 2015). Briefly, the cerebellar cortex was removed and dissociated with papain, granule neuron progenitors were purified by centrifugation through a Percoll gradient, and neurons were plated on poly-D-lysine coated plates in neurobasal media with B27 supplements (Invitrogen), 1% FBS, and pen-strep.

On day 1 of *in vitro* culture (DIV1), neurons were transduced with lentiviruses. All neurons received either 2 gRNAs targeting the *Grin2c* promoter (Site 2-2. Site 2-3 from Frank et al. (Frank et al., 2015) or a control non-targeting gRNA complementary to a sequence in the LacZ gene (Chen et al., 2019). gRNAs were delivered in a FUGW-based U6 chimeric gRNA expression vector co-expressing GFP. In addition to the gRNA virus, one set of cells received a lentivirus expressing the Cre recombinase under the control of the hSyn1 promoter (Addgene 86641). This induced expression of the dCas9^p300^ transgene in the neurons. Other neurons received lentivirus expressing dCas9^VP64^ (Perez-Pinera et al., 2013) with the gRNAs to compare the functions of the two transgenes in neurons from the same cultures. Viruses were titered on HEK293T cells before use and delivered at a MOI of 10.

### AAV cloning and production

An AAV backbone containing a gRNA cloning site, pCBh.Cre-WPRE-hGHpA was obtained from Addgene (plasmid #60229). Two oligonucleotides per gRNA were purchased, annealed, and cloned into the SapI cloning site. Sequences for protospacers can be found in Supplementary Table 1. Prior to AAV production, ITRs were verified by SmaI digest. pCre-*Pdx1.gRNA* and pCre-control.gRNA were used to generate high titer AAV9. Viral supernatants at the indicated titers were dialyzed w/350 mM NaCl and 5% D-sorbitol in PBS. For neurons, gRNAs for LacZ or *Fos* Enh2 sequences were cloned in the Sap1 site of an AAV backbone (Addgene #60231) with hSyn-Cre-2A-GFP. High titer AAVs for intracranial delivery were produced in serotype AAV2/1 by the Duke University Viral Vector Core Facility.

### Lentiviral cloning and production

Lentiviral production was performed using standard procedures in HEK293T cells using second generation lentiviral vectors VSVg and d8.9 (Salmon and Trono, 2006). Lentiviral preps for fibroblast transduction were purified using Lenti-XTM (Clonetech/Cat#631232) to a 20X concentration. Fibroblasts were transduced at 1X concentration in the absence of polybrene for 24 hours, media was then changed and fibroblasts were returned to growth media. For neurons, viruses were purified by ultracentrifugation and resuspended at a titer of ~5×10^5^ infectious particles per μL of PBS. Neurons were transduced in a solution of DMEM + 0.5 μg/mL polybrene for 6 hrs, then neurons were washed and returned to conditioned medium for 5 days to allow for viral gene expression.

### RNA isolation and qRT-PCR

RNA was isolated from fibroblasts using a Qiagen RNAeasy (Cat# 74136). cDNA synthesis was performed using Superscript III with Vilo buffer. For *Pdx1* quantification, Taqman probes (*Pdx1* (4453320 (Mm00435565_m1) and *Gapdh* (4448489 (Mm99999915_g1) were used along with Perfecta qPCR Fastmix II (Cat #95118-012). For *Pcsk9* quantification, primers and conditions from a previously published report were used (Thakore et al., 2018).

Neurons were harvested for RNA extraction (Qiagen RNAeasy kit with on-column DNAse treatment), oligo(dT) and random primers primed for cDNA synthesis with iScript™ cDNA Synthesis Kit (BioRad Cat#1708891), and run for SYBR qPCR on a Quantstudio 3 realtime PCR system (Thermo). All gene expression values were normalized to *Gapdh* in the same well to control for sample handling. Sequences for qPCR primers can be found in Supplemental Table 2.

T cells were collected in Cells-to-CT 2-Step Taqman kit (Cat# A35377) following the manufacturer’s instructions. *Foxp3* quantification was performed using Taqman probes *Foxp3*(4331182(Mm00475162_m1)) and *Gapdh* (4448489 (Mm99999915_g1)

### RNA-seq and differential expression analysis

For liver tissue, RNA was isolated using the Qiagen RNA plus universal Kit (Cat# 73404). RNA-seq was performed after Total RNA was polyA tail enriched, rRNA depleted, and libraries prepped with a NEBNext Ultra II kit (E7645L). For T cells, transduced cells were FACS purified using an SH800 and fluorescent antibody targeting Thy1.1 that was expressed from the same vector as the gRNA cassette. Cell pellets were processed using Qiagen RNeasy RNA extraction kit. RNA-seq libraries were built from rRNA depleted mRNA using the Illumina TruSeq mRNA library prep kit (Cat# 20020595). All RNA-seq samples were first validated for consistent quality using FastQC v0.11.2 (Babraham Institute). Raw reads were trimmed to remove adapters and bases with average quality score (*Q*) (Phred33) of < 20 using a 4bp sliding window (SLIDINGWINDOW:4:20) with Trimmomatic v0.32 (Bolger et al., 2014). Trimmed reads were subsequently aligned to the primary assembly of the GRCm38 mouse genome using STAR v2.4.1a (Dobin et al., 2013) removing alignments containing non-canonical splice junctions (--outFilterIntronMotifs RemoveNoncanonical).Aligned reads were assigned to genes in the GENCODE vM13 comprehensive gene annotation (Harrow et al., 2012) using the featureCounts command in the subread package with default settings (v1.4.6-p4 (Liao et al., 2013)). Differential expression analysis was performed using DESeq2 (v1.22.0 (Love et al., 2014)) running on R (v3.5.1). Briefly, raw counts were imported and filtered to remove genes with low or no expression, keeping genes having two or more counts per million (CPMs) in two or more samples. Filtered counts were then normalized using the DESeq function which uses estimated size factors to account for library size as well as gene and global dispersion. In order to find significant differentially expressed genes, the nbinomWaldTest was used to test the coefficients in the fitted Negative Binomial GLM using the previously calculated size factors and dispersion estimates. Genes having a Benjamini-Hochberg false discovery rate (FDR) less than 0.05 were considered significant (unless otherwise indicated). Log2 fold-change values were shrunk towards zero using the adaptive shrinkage estimator from the ‘ashr’ R package (Stephens, 2017). For estimating transcript abundance, Transcript Per Million (TPMs) were computed using the rsem-calculate-expression function in the RSEM v1.2.21 package (Li and Dewey, 2011). Data were visualized using pandas v0.23.3 and seaborn v0.9.0 packages in Python 2.7.11.

Raw sequencing files are available from the NCBI Gene Expression Omnibus via SuperSeries accession GSE146848.

### Protein isolation and western blot

Protein from tissue was isolated using a Biomasher II mortar and pestle tubes in conjunction with RIPA or TGH buffer. Protein was quantified using BCA assay. Novex NuPage 4-12% Bis-Tris gels were run for 45 minutes in MES running buffer and transferred onto nitrocellulose membranes for an hour at 4 °C in towbin’s buffer with 20% MeOH. Membranes were blocked for non-specific binding overnight in 5% milk in TBS-T. 1° Antibodies were incubated in 5% milk in TBS-T and used at the following manner: Anti-Cas9 (EnCor Biotechnology/Cat#MCA-3F)-1:2000 at room temp for 2-3 hours, Anti-Gapdh (Cell Signaling/Cat#2118L) 1:5000, Anti-LDLR (Abcam/Cat# ab52818)-1:1000, and anti-Actin (EMD Millipore/Cat#MAB1501) overnight at 4°C. After washing, secondary antibodies (Anti-rabbit HRP (Sigma/Cat#A6154) and Anti-Mouse HRP (Santa Cruz/Cat#sc-2005) were incubated in 3% BSA in TBS-T at 1:5000. Membranes were visualized using Clarity western ECL substrate (Cat#1705061) from BioRad and imaged using a BioRad ChemiDoc XRS+. For neurons, cells were lysed directly in 1X RIPA buffer and blots were incubated with goat anti-mouse 680 (1:5000, cat #20253, Biotium). Fluorescent immunoreactivity was imaged on a LICOR Odyssey.

### Stereotaxic injection of gRNA AAVs in mouse hippocampus

AAV containing GFP and a gRNA targeting either LacZ (control) or *Fos* Enh2 (experimental treatment) were stereotaxically injected into each hemisphere of the dorsal hippocampus of adult male dCas9^p300^ heterozygous mice such that each mouse had one control side and one experimental side (stereotaxic coordinates AP: −2.3, ML +/−1.8, DV: −1.8). 3 wks following transduction, mice were either used for evaluation of Fos expression (n=3 mice) or for electrophysiology (n=2 mice).

### Histological staining

For PDX1 staining, livers were fixed overnight with 4% PFA and 12 μm-thick cryostat sections were processed for immunofluorescence. Slides were washed in PBS (Phosphate-buffered saline), boiled in citrate buffer (10 mM citric acid, 0.05% Tween-20, pH 6.0) for 25 min at 95 °C and were then kept for 20 minutes at room temperature (antigen retrieval). Slides were permeabilized with PBS containing 0.2% Triton for 20 minutes, washed with PBS and blocked with 3% BSA + 0.1% Tween in PBS for 1 hour. Primary antibody (Abcam, ab47267, 1:100) was incubated overnight at 4 °C in blocking reagent. Sections were then washed and incubated with specific secondary antibody coupled to Alexa-Fluor-488 (Life Technologies, 1:200) and DAPI. Confocal images were acquired with a Zeiss LSM 880 microscope.

For Fos staining in hippocampal sections, after 3 weeks following viral injection, mice were placed in an open field and allowed to explore three novel objects for a period of 2 hours. Mice were then perfused with 4% paraformaldehyde and brains were coronally sectioned on a freezing microtome for immunostaining. Primary antibodies used were rabbit anti-c-Fos (Calbiochem #PC38, 1:1000) or rabbit anti-Cas9 (EnCor Biotechnology, Cat# RPCA-CAS9-Sp, 1:1000). These were detected by anti-rabbit Cy3 (1:500). Z-stack images through the dentate gyrus were obtained using a Leica SP8 upright confocal with a 40X objective plus additional digital zoom. For each animal, the hemisphere infected with gRNA targeting *Fos* Enh2 was compared to the control hemisphere expressing the control LacZ gRNA. To count high Fos-expressing cells, z-stacks were converted to sum projections and thresholded using ImageJ. These counts were compared using a paired t-test. To determine whether there were changes in Fos intensity across the population of GFP-positive cells, ROIs were created for each GFP-positive cell within a given region while blind to Fos expression. Fluorescence intensity of the Fos channel was then measured for each of these cells, creating a distribution of Fos intensities. The Fos signal for each cell was normalized to the average control hemisphere Fos value for a given animal. The resulting distributions were then compared using a Kolmogorov-Smirnoff test.

### ChIP-Sequencing and analysis

Livers were pulverized and fixed as described by Savic et al (Savic et al., 2013). Following fixation, livers were rocked at 4°C in 10 ml Farnham Lysis Buffer for 10 minutes followed by centrifugation. Pellets were resuspended in 4 mL RIPA buffer. 1 mL RIPA and 1 mL RIPA containing resuspended liver was placed into a 15 mL tube with 800 mg of Diagenode sonication beads (Diagenode, C01020031). This tube was sonicated for 30 cycles of 30s on and 30s off using a Bioruptor Pico. ChIP was performed using Cas9 (Diagenode C15200229-100), Flag-M2 (Sigma, F1804), H3K27ac (Abcam, ab4729), or H3K9me3 (Abcam, ab8898) antibodies. Following ChIP, ChIP-seq libraries were prepared using a Kapa Hyper Prep kit and libraries sequenced using 50bp single end reads on a Illumina hiseq4000. For analysis, adapter sequences were removed from the raw reads using Trimmomatic v0.32 (Bolger et al., 2014). Reads were aligned using Bowtie v1.0.0 (Langmead et al., 2009), reporting the best alignment with up to 2 mismatches (parameters --best --strata -v 2). Duplicates were marked using Picard MarkDuplicates v1.130 (http://broadinstitute.github.io/picard/), while low mappability or blacklisted regions identified by the ENCODE project were filtered out from the final BAM files. Signal files were generated with deeptools bamCoverage (v3.0.1 (Ramirez et al., 2014)) ignoring duplicates, extending reads 200bp and applying RPKM normalization. Using the sequenced input controls, binding regions were identified using the callpeak function in MACS2 v2.1.1.20160309 (Zhang et al., 2008). For the differential binding analysis, first, a union peakset was computed merging individual peak calls using bedtools2 v2.25.0 (Quinlan and Hall, 2010). Then, reads in peaks were estimated using featureCounts from the subread package v1.4.6-p4 (Liao et al., 2013). The difference in binding was assessed with DESeq2 v1.22.0 (Love et al., 2014) using the nbinomWaldTest to test coefficients in the fitted Negative Binomial GLM. Data were visualized using pandas v0.23.3 and seaborn v0.9.0 packages in Python 2.7.11 or R 3.6.5.

For cerebellar granule neurons, chromatin immunoprecipitation was performed following the protocol of EZ-ChIP (Millipore, 17-371). Briefly, cells were lysed by SDS Lysis Buffer and sonicated for 1.5 hrs (Diagenode Bioruptor) at 4°C on the high setting with 30 seconds on/off interval. 20 μl Dynabeads Protein G (ThermoFisher, 10003D) was pre-incubated with 2 μg antibodies in 1XPBS buffer for 4 to 6 hours at 4°C. Mouse anti-Cas9 (Encor MCA-3F9 and Diagenode C15200229-100) antibodies were used. Cell lysates were then incubated overnight with antibody-bead complexes at 4°C. Subsequently, the beads were washed with Low Salt Immune Complex Wash Buffer, High Salt Immune Complex Wash Buffer, LiCl Immune Complex Wash Buffer, and TE Buffer. Bound protein/DNA complexes were eluted by ChIP elution buffer and then reversed the crosslinks. Samples were treated with RNase A and Proteinase K for post-immunoprecipitation and then the DNA was purified using QIAquick PCR Purification Kit (Qiagen, 28104). ChIP primers sequences can be found in Supplemental Table 3. ChIP for *Grin2c* was normalized to ChIP for *Gapdh* in the same sample to control for concentration and sample handling.

For CD4+ Tcells, 2 million cells were fixed with 1% paraformaldehyde at room temperature for 10 minutes, quenched with 0.125M glycine and washed twice with PBS. Fixed cells were lysed with 1mL of Farnhyme Lysis Buffer for 5 minutes on ice followed by centrifugation at 800 x g for 5 minutes at 4°C. Cell pellets were snap frozen before being resuspended in 100μL of RIPA buffer. Cells were then sonicated for 10 cycles of 30 seconds on/off on a Bioruptor Pico (Diagenode) at 4°C. Chromatin immunoprecipitation and library prep was performed with the anti-H3K27ac (Abcam, ab4729) antibody as previously described (Ref GGR)

Raw sequencing files are available from the NCBI Gene Expression Omnibus via SuperSeries accession GSE146848.

### Hippocampal slice electrophysiological recordings

3 wks after viral injections into dCas9^p300^ heterozygous mice, mice were anesthetized with isofluorane, and transcardially perfused with ice-cold NMDG artificial cerebrospinal fluid (NMDG-ACSF; containing 92 mM NMDG, 2.5 mM KCl, 1.2 mM NaH2PO4, 30 mM NaHCO3, 20 mM HEPES, 2 mM glucose, 5 mM sodium ascorbate, 2 mM thiourea, 3 mM sodium pyruvate, 10 mM MgSO4, 0.5 mM CaCl2) that was bubbled with 5% CO2/95% O2. The brain was extracted and sectioned into 300 μm thick sagittal slices using a vibratome (VT-1000S, Leica Microsystems) in ice-cold oxygenated NMDG-ACSF. Coronal sections including the dorsal hippocampus were bubbled in the same solution at 37 °C for 8 min, and transferred to oxygenated modified-HEPES ACSF at room temperature (20–25 °C; 92 mM NaCl, 2.5 mM KCl, 1.2 mM NaH2PO4, 30 mM NaHCO3, 20 mM HEPES, 2 mM glucose, 5 mM sodium ascorbate, 2 mM thiourea, 3 mM sodium pyruvate, 2 mM MgSO4, 2 mM CaCl2) for at least 1 h before recording. Recordings were performed in a submerged chamber, superfused with continuously bubbled ACSF (125 mM NaCl, 2.5 mM KCl, 1.25 mM NaH2PO4, 25 mM NaHCO3, 25 mM glucose, 2 mM CaCl2, 1 mM MgCl2, 2 mM sodium pyruvate) at near-physiological temperature (34 ± 1 °C). For whole-cell current-clamp recordings, the electrodes (4–6 MΩ) were filled with an intracellular solution containing 130 mM K-methylsulfonate, 5 mM NaCl, 10 mM HEPES, 15 mM EGTA, 12 mM phosphocreatine, 3 mM Mg-ATP, 0.2 mM Mg-GTP, 0.05 AlexaFlour 594 cadaverine, and 1% biotin. Current clamp responses were recorded with a Multiclamp 700B amplifier (filtered at 2-4 kHz) and digitized at 10 kHz (Digidata 1440). Series resistance was always <20 MΩ and was compensated at 80%-95%. Virally transduced pyramidal cells expressing GFP were visualized by infrared differential interference contrast and fluorescence video microscopy (Olympus; CoolLED) with a 40x objective. Data were collected and analyzed off-line using AxographX and Neuromatic package (Think Random) in Igor Pro software (WaveMetrics). Resting membrane potential was measured while cells were held at 0 pA. The action potential frequency was measured during holding currents of 0 to 500 pA. Rheobase was measured as the minimal amount of current needed to depolarize the cell membrane. Initial spike latency was defined as the time from the onset of current holding change to the first measurable deflection of the potential from baseline. Membrane time constant was defined as the time it took the cell’s voltage level to decay to resting state after delivering a 200ms step of −150pA. Input resistance was measured using “∆V/pA =mΩ” for each current step and averaging across all current steps for individual neurons. All measurements were averaged across neurons. Finally, recorded neurons were labeled with biotin delivered through the patch pipette, and post-hoc confocal imaging of fixed slices confirmed GFP expression in the recorded neurons.

### Novel object exploration and hippocampal expression of Fos

AAV vectors containing GFP and gRNA targeting either LacZ (control) or *Fos* Enh2 were injected into either hemisphere of the dorsal hippocampus of adult male dCas9^p300^ mice such that each mouse had one control hemisphere and one experimental hemisphere (stereotaxic coordinates AP: −2.3, ML +/−1.8, DV: −1.8). Three weeks following AAV injection, mice were placed in an open field and allowed to explore three novel objects for a period of 2 hours. Mice were then perfused with 4% paraformaldehyde and brains were coronally sectioned on a freezing microtome for immunostaining. Primary antibodies used were rabbit anti-c-Fos (Calbiochem #PC38, 1:1000) and rabbit anti-Cre recombinase (Cell Signaling #15036, 1:1000). These were detected by anti-rabbit Cy3 (1:500). Z-stack images through the dentate gyrus were obtained using a Leica SP8 upright confocal with a 40X objective plus additional digital zoom. For each animal, the hemisphere infected with gRNA targeting *Fos* Enh2 was compared to the control hemisphere expressing the control LacZ gRNA. To count high Fos-expressing cells, z-stacks were converted to sum projections and thresholded using ImageJ. These counts were compared using a paired t-test. To determine whether there were changes in Fos intensity across the population of GFP-positive cells, ROIs were created for each GFP-positive cell within a given region while blind to Fos expression. Fluorescence intensity of the Fos channel was then measured for each of these cells, creating a distribution of Fos intensities. The Fos signal for each cell was normalized to the average control hemisphere Fos value for a given animal. The resulting distributions were then compared using a Kolmogorov-Smirnoff test.

### Purification of CD4 T cells for in vitro cultures

Spleens and lymphnodes were harvested, dissociated, and processed with ammonium chloride potassium (ACK) lysis buffer and passed through the Magnisort mouse CD4 T cell enrichment kit (Thermo cat# 8804-6821-74). Lymphocytes were surface-stained with antibodies for the purification of nT cells (CD4^+^CD25^−^CD62L^hi^CD44^lo^) through FACS using either a Beckman Culture Astiros (CD4-FITC; CD25-eFluor450; CD62L-APC; CD44-PE, and fixable viability dye e780) or SONY SH800 (CD4-PECy7; CD25-PE; CD44-FITC; CD62L-PEcy5; Live/Dead Red) instrument to achieve ≥98% purity. For ChIP-seq, Tn cells were purified using Magnisort mouse Naïve CD4+ Tcell enrichment kit (Thermo cat# 8804-6824-74) rather than sorting in order to increase starting material. Antibody information: CD4 (eBiosciences, Cat #25-0042-82), CD25 (eBiosciences, Cat# 12-0251-83), CD44 (eBiosciences Cat# 11-0441-85), CD62L (eBiosciences, Cat# 15-0621-82).

### *In vitro* culture conditions for CD4 T cell subset polarization

All T cell polarizations were performed using Olympus tissue-culture plastic that was coated with anti-Hamster IgG (20ng/μL, MP Biomedical Cat# 856984) at 37°C for at least 4 hours prior. Tn cells were seeded at ~250k cells/mL. Th0 activating conditions contained hamster-anti-CD3e (0.25 μg/mL), hamster-anti-CD28 (1 μg/mL), blocking antibodies anti-IL4 (2μg/mL, eBiosciences Cat# 16-7041-85) and anti-IFNg (2μg/mL, eBiosciences Cat# 16-7311-85), and mIL-2 (10 ng/mL, peprotech Cat# 212-12). iTreg polarizing conditions contained the above and was supplemented with hTGFb1 (5 ng/mL, eBiosciences Cat# 14-8348-62).

### Treg effector generation and spinfection

Tn cells were isolated and purified from the spleen and lymph nodes of *Cd4:Cre+/dCas9-p300+/Foxp3-eGFP+* mice via FACS as described above and cultured in Th0 activating conditions at 37°C and 5% CO_2_. Between 20-24 hours later, the activated T cells were transduced via spinfection with viral supernatant containing 6.66 ng/mL polybrene and Mouse Stem Cell Virus (MSCV) harboring a gRNA expression vector with either a single gRNA targeting the *Foxp3* promoter, or a non-targeting control gRNA. Viral supernatant was replaced with Th0 or iTreg media and cells were cultured for 3 or 7 days.

### Purification of effector T cells for co-culture

At 7 days after viral transduction, Treg effector cells were stained with fixable viability dye e780 (eBioscience Cat# 65-0865-18), Thy1.1 (CD90.1-PE, eBiosciences, Cat# 45-0900-82). Viable transduced cells expressing Foxp3-eGFP were FACS purified (Thy1.1^+^/FOXP3-eGFP+) and resuspended at 500,000 cells/mL in RPMI.

### Purification of effector T cells for ChIP and RNA-seq

Tn cells were activated for 20-24 hours in Th0 conditions as above before transduction with *Foxp3* or control gRNA. 3 days post transduction, 100K or 2M viable, Thy1.1^+^ transduced Tcells were FACS purified for RNA-seq or ChIP-seq, respectively.

### Purification of conventional T cells and suppression assay

Spleens and lymphnodes from *Foxp3*-eGFP mice (JAX stock# 006769) were processed as above and the recovered splenocytes were stained for viability (Live/Dead red; Thermo, cat# L34971) and CD4-PEcy7. CD4^+^FOXP3^−^ live cells were purified using a SONY SH800 FACS machine. Cells were washed with 30mL of PBS and stained with 5 μM Cell Trace Violet (CTV; Thermo Cat# C34557) following manufacturer’s instructions. Stained Tconv cells were diluted to 500,000 cells/mL in RPMI and co-cultured with 25,000 Treg cells per well for 72 hours with 25,000 aCD3/aCD28 Dynabeads (Thermo cat# 11161). Cells were then removed from the magnetic beads, stained for viability with e780, and processed for flow cytometric analysis.

### FOXP3 protein staining and flow cytometry analysis

T cells were stained with Thy1.1-PerCP-Cy5.5 (eBioscience cat # 45-0900-82) and fixable viability dye eFluor780 prior to fixation and permeabilization with FOXP3 Transcription Factor Staining Buffer Kit (eBioscience cat# 00-5523-00) following manufacturer’s protocol. Cells were stained intracellularly with Foxp3-PE (eBioscience cat#12-5773-82) and analyzed with a BD FACSCanto II cytometer.

