## Supplementary Figures for "Transgenic mice for *in vivo* epigenome editing with CRISPR-based systems"

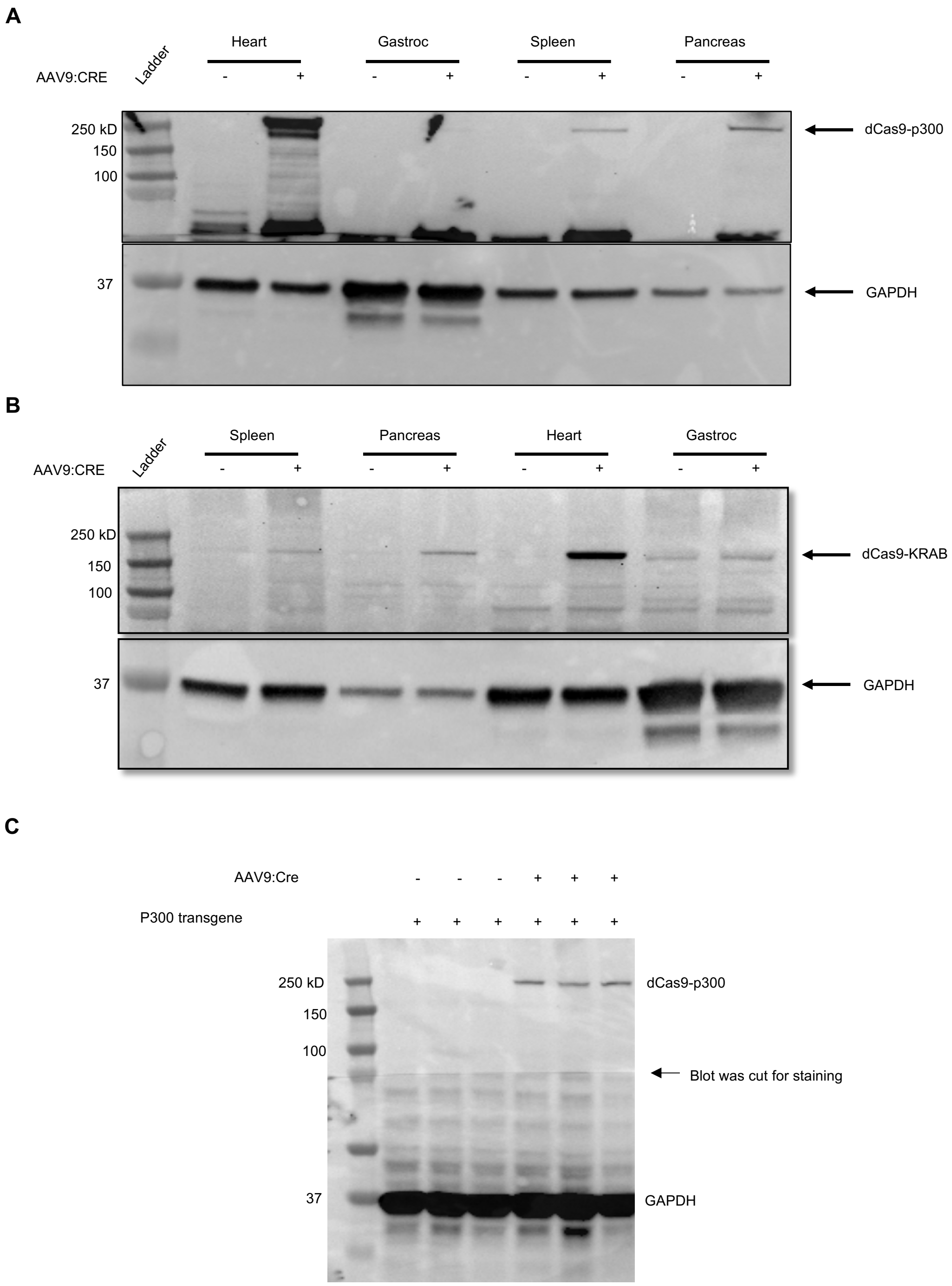


**Supplemental Figure 1**: **Western blot analysis of dCas9$^{p300}$ or dCas9$^{KRAB}$ protein levels following systemic administration of AAV9:CMV.Cre**. A) High levels of dCAS9$^{p300}$ are detected 14 days post injection in heart, moderate levels are seen in the spleen and pancreas, and low levels in the gastrocnemius. B) Similarly, high levels of dCAS9$^{KRAB}$ is found in the heart while moderate levels are seen in the spleen and pancreas at 8 weeks post injection. C) dCAS9$^{p300}$ protein levels in the liver of AAV9:CMV.Cre treated animals 14 days post-treatment with or without *Cre*.

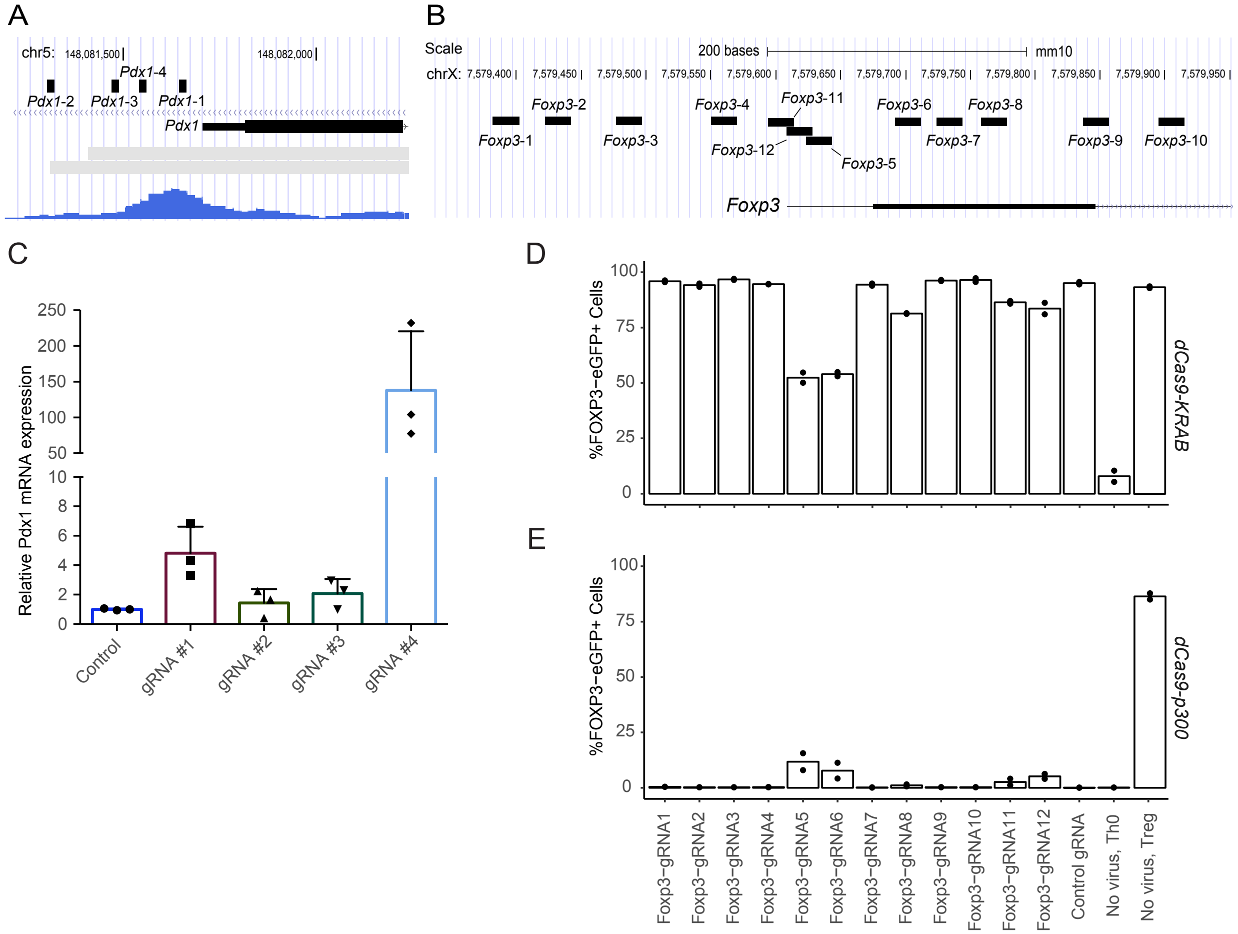


**Supplemental Figure 2: Testing candidate gRNAs in primary fibroblasts and T cells from *Rosa26:LSL-dCas9$^{p300}$* mice.** A) Genome browser track showing the location of the four tested gRNAs for the *Pdx1* promoter region and B) twelve gRNAs for the *Foxp3* promoter region. C) *Pdx1*-targeting gRNAs were tested in primary fibroblasts from the hindlimb of a *Rosa26:LSL-dCas9$^{p300}$* mouse in triplicate and compared to a *Myod*-targeting control gRNA. gRNA #4 significantly upregulated *Pdx1* as compared to controls. (n=3 per gRNA, ANOVA with Dunnett's post-hoc, $p < 0.0035$). D) Percentage of FOXP3-eGFP positive cells after dCas9$^{p300}$-induced activation with *Foxp3*-targeting gRNAs in Th0 cells from a *Rosa26:LSL-dCas9$^{p300}$* mouse and E) dCas9$^{KRAB}$-induced repression with *Foxp3*-targeting gRNAs in Treg cells from a *Rosa26:LSL-dCas9$^{KRAB}$* mouse compared to a control non-targeting gRNA and non-transduced Th0 or Treg cells.

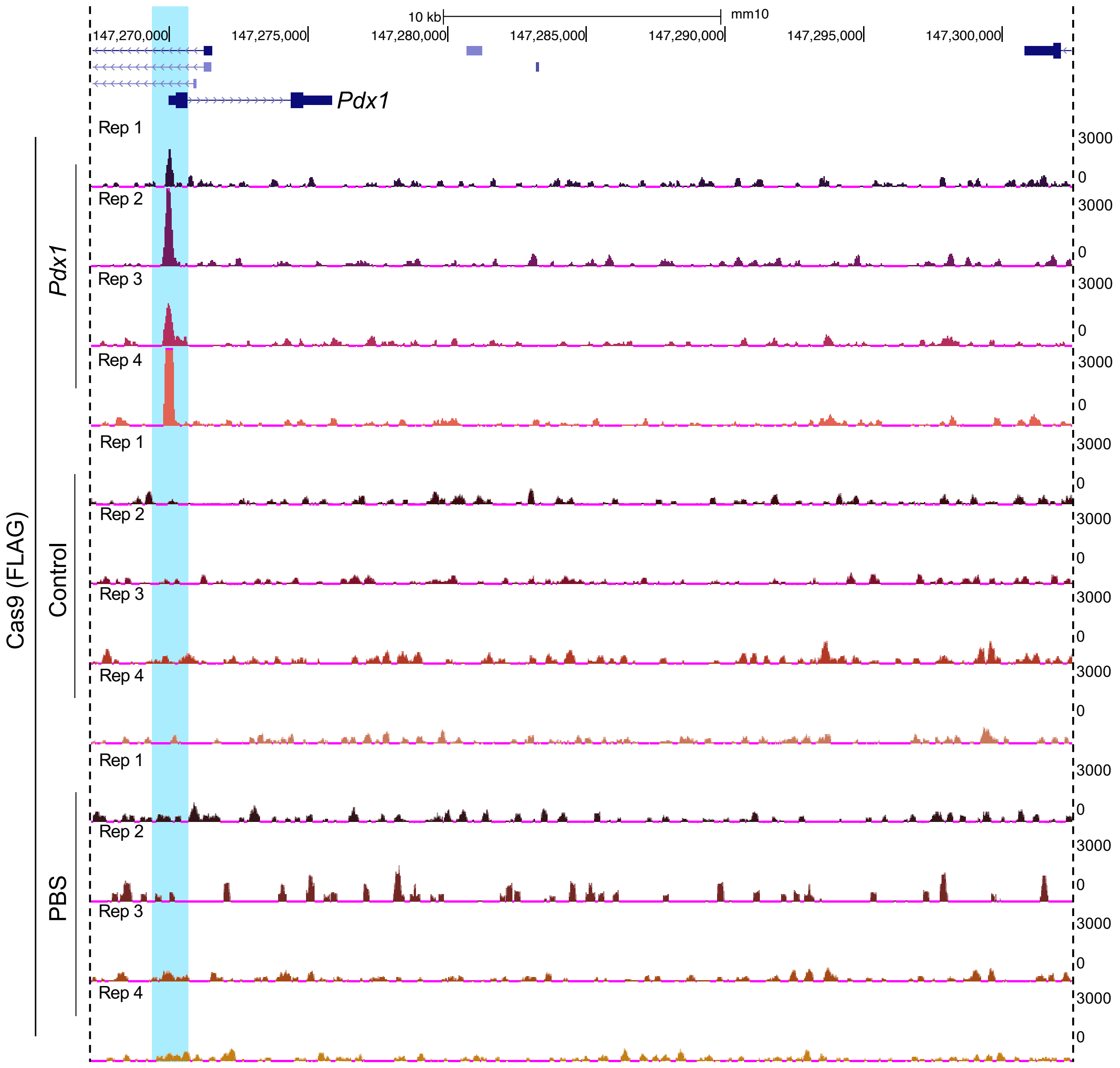


**Supplemental Figure 3: Specificity of dCas9$^{p300}$ binding the *Pdx1* promoter *in vivo* in the liver of *Rosa26:LSL-dCas9$^{p300}$* mice following systemic administration of AAV9 encoding both *Cre* and a gRNA.** Genome browser tracks showing Cas9 ChIP-seq signal from livers of dCas9$^{p300}$ mice 14 days post-treatment with an AAV9:Cbh.Cre-gRNA, either *Pdx1*-gRNA or the control-gRNA, and mice treated with saline (PBS). The highlighted region shows the area where dCas9$^{p300}$ is targeted to the promoter of *Pdx1*. Specific signal is found only in the *Pdx1*-gRNA treated animals.

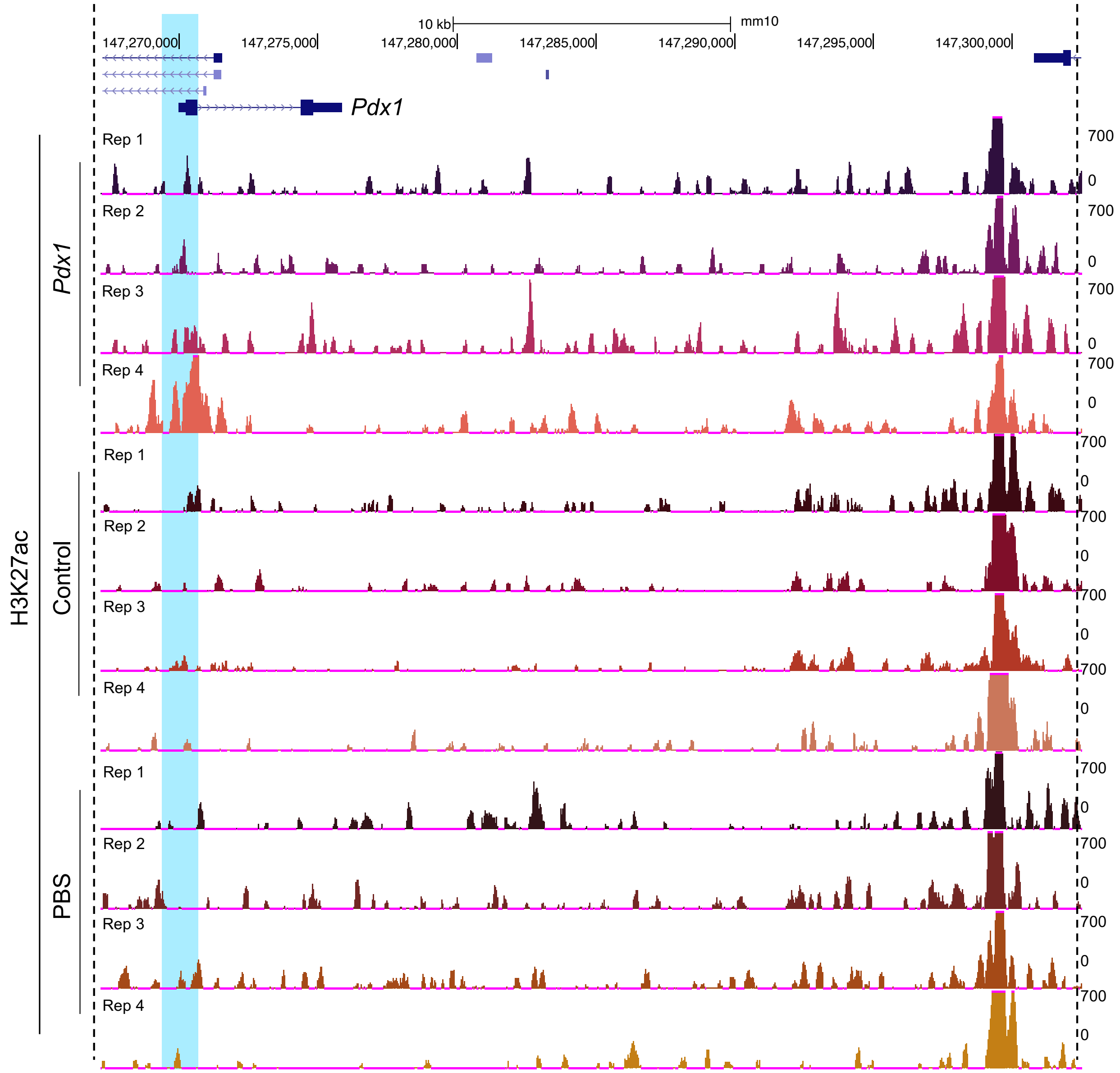


**Supplemental Figure 4: H3K27ac enrichment at the *Pdx1* promoter *in vivo* in the liver of *Rosa26:LSL-dCas9$^{p300}$* mice following systemic administration of AAV9 encoding both *Cre* and a gRNA.** Genome browser tracks of H3K27ac ChIP-seq signal from livers of dCas9$^{p300}$ mice 14 days post-treatment with AAV9:Cbh.Cre-gRNA, encoding either the *Pdx1*-targeting gRNA or the control non-targeting gRNA, and mice treated with saline (PBS). The highlighted region shows the area where dCas9$^{p300}$ is targeted to the promoter of *Pdx1*. Enrichment of H3K27ac around the dCAS9$^{p300}$ target site in the *Pdx1* gRNA treated mice is highlighted in blue.

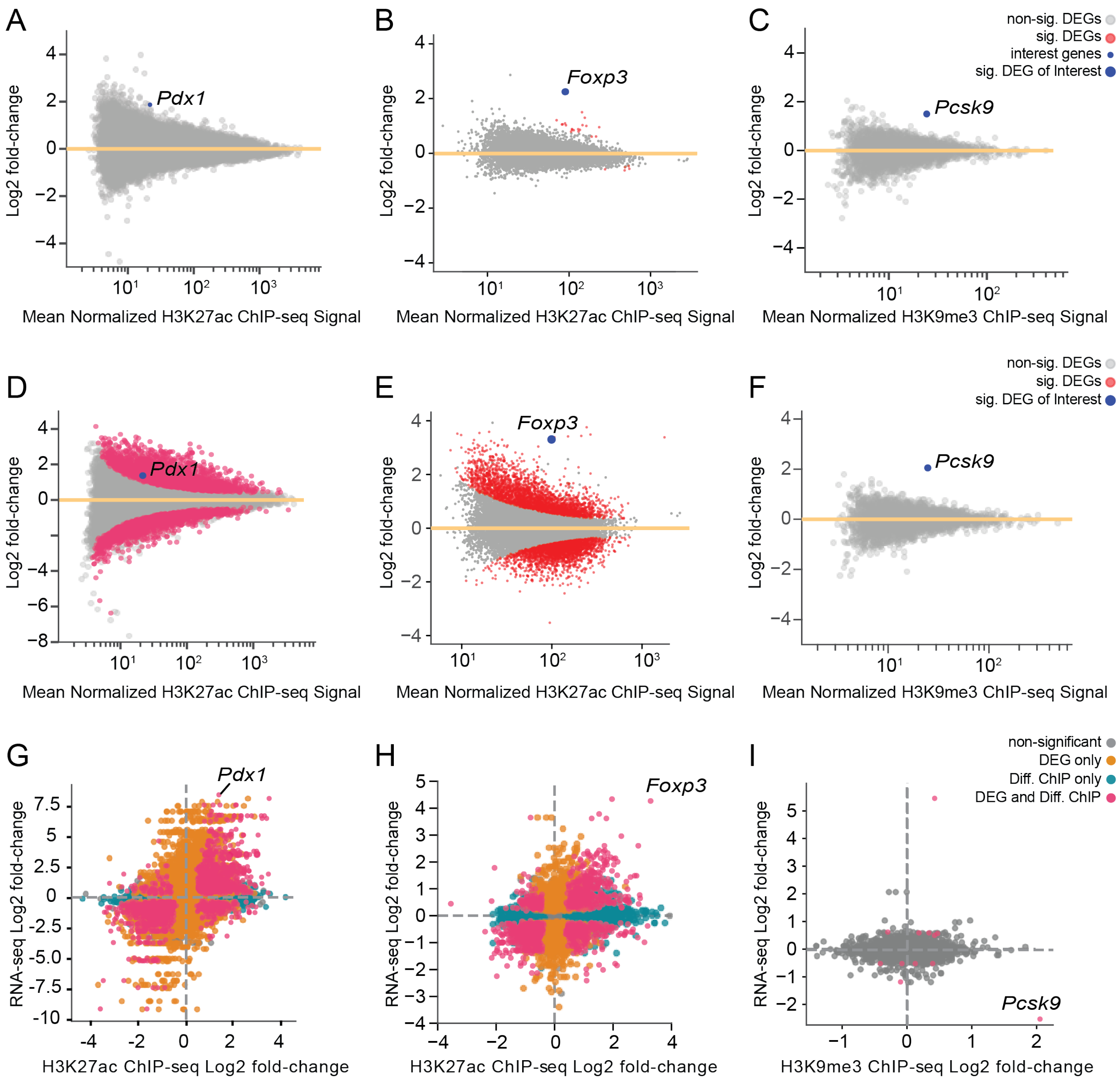


**Supplemental Figure 5: Genome-wide analysis of histone ChIP-seq data from *Rosa26:LSL-dCas9*$^{p300}$ and *dCas9*$^{KRAB}$ mice in liver samples and T cells after transduction.** A) MA plots show $\log_2$(fold-change) in H3K27ac enrichment for samples from *Rosa26:LSL-dCas9*$^{p300}$ mice when comparing A) liver cells treated with *Pdx1*-targeting gRNA vs control non-targeting gRNA, B) Th0 cells from mice crossed to *Cd4:Cre* and treated with *Foxp3*-targeting gRNA vs control non-targeting gRNA, and C) H3K9me3 enrichment from *Rosa26:LSL-dCas9*$^{KRAB}$ mouse liver samples treated with *Pcsk9*-targeting gRNA vs control non-targeting gRNA. $\text{Log}_2$(fold-change) in D) H3K27ac enrichment in samples from *Rosa26:LSL-dCas9*$^{p300}$ mice when comparing liver cells treated with *Pdx1*-targeting gRNA vs saline, E) H3K27ac enrichment in Th0 cells from *Rosa26:LSL-dCas9*$^{p300}$ mice treated with *Foxp3*-targeting gRNA with vs without *Cd4:Cre*, and F) H3K9me3 enrichment from *Rosa26:LSL-dCas9*$^{KRAB}$ mouse liver samples treated with *Pcsk9*-

targeting gRNA vs saline. G) Relationship of $\log_2$(fold-change) in read counts per million (CPM) from RNA-seq and H3K27ac ChIP-seq when comparing between *Rosa26:LSL-dCas9*$^{p300}$ mouse liver samples treated with Pdx1-targeting gRNA vs saline, H) *Rosa26:LSL-dCas9*$^{p300}$ Th0 cells treated with *Foxp3*-gRNA with or without *Cd4:Cre*, and I) the relationship between RNA-seq and H3K9me3 ChIP-seq when comparing *Rosa26:LSL-dCas9*$^{KRAB}$ mouse liver samples treated with *Pcsk9*-targeting gRNA vs saline control (all data are N=3 or 4 mice per treatment group; orange = significant differentially expressed gene (DEG) only; blue = significant differently enriched ChIP-seq signal only; red significant differently expressed gene and ChIP-seq enrichment; grey = non-significant locus, FDR < 0.05 for liver samples, FDR < 0.01 for T cell samples)

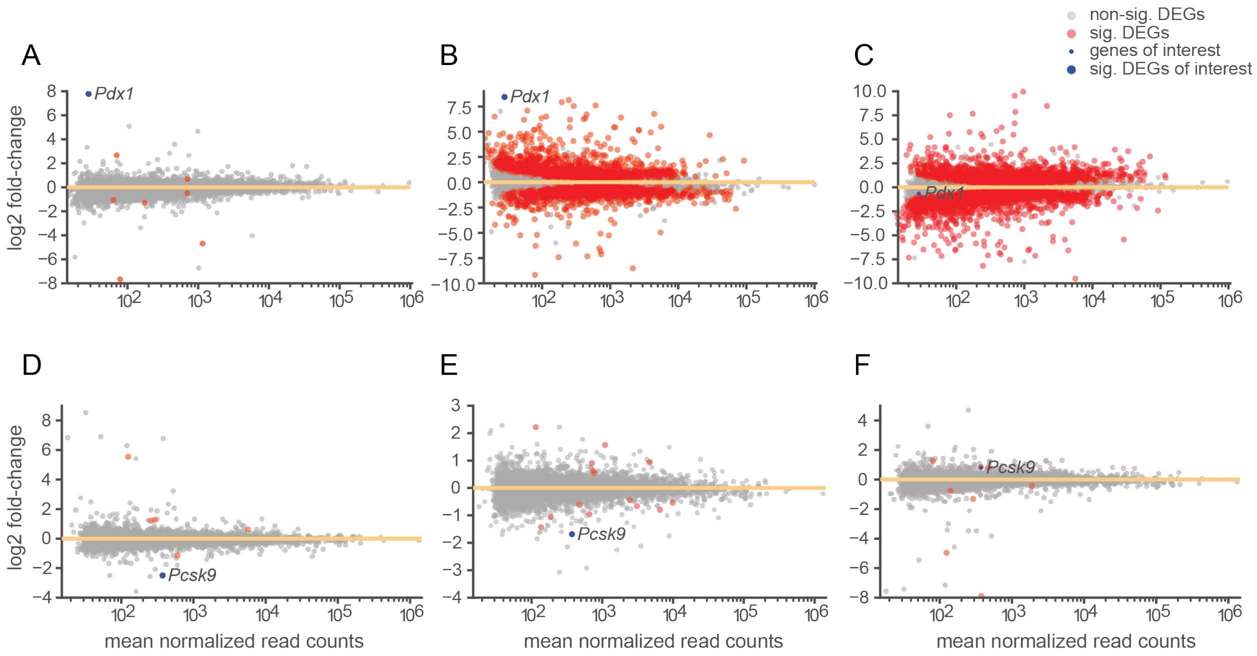


**Supplemental Figure 6: Differential analysis of RNA-seq data from dCas9 epigenome editor mouse liver samples after transduction.** A) MA plots show $\log_2$(fold-change) in gene expression for *Rosa26:LSL-dCas9*$^{p300}$ mouse liver when comparing between *Pdx1*-targeting gRNA vs control non-targeting gRNA, B) *Pdx1*-targeting gRNA vs saline, and C) control non-targeting gRNA vs saline. D) $\text{Log}_2$(fold-change) in gene expression for *Rosa26:LSL-dCas9*$^{KRAB}$ mouse liver when comparing between *Pcsk9*-targeting gRNA vs control non-targeting gRNA, E) *Pcsk9*-targeting gRNA vs PBS, and F) control non-targeting gRNA vs PBS. Red and blue dots indicate significant changes in gene expression. All data are average of N=4 mice per treatment group, FDR < 0.05.

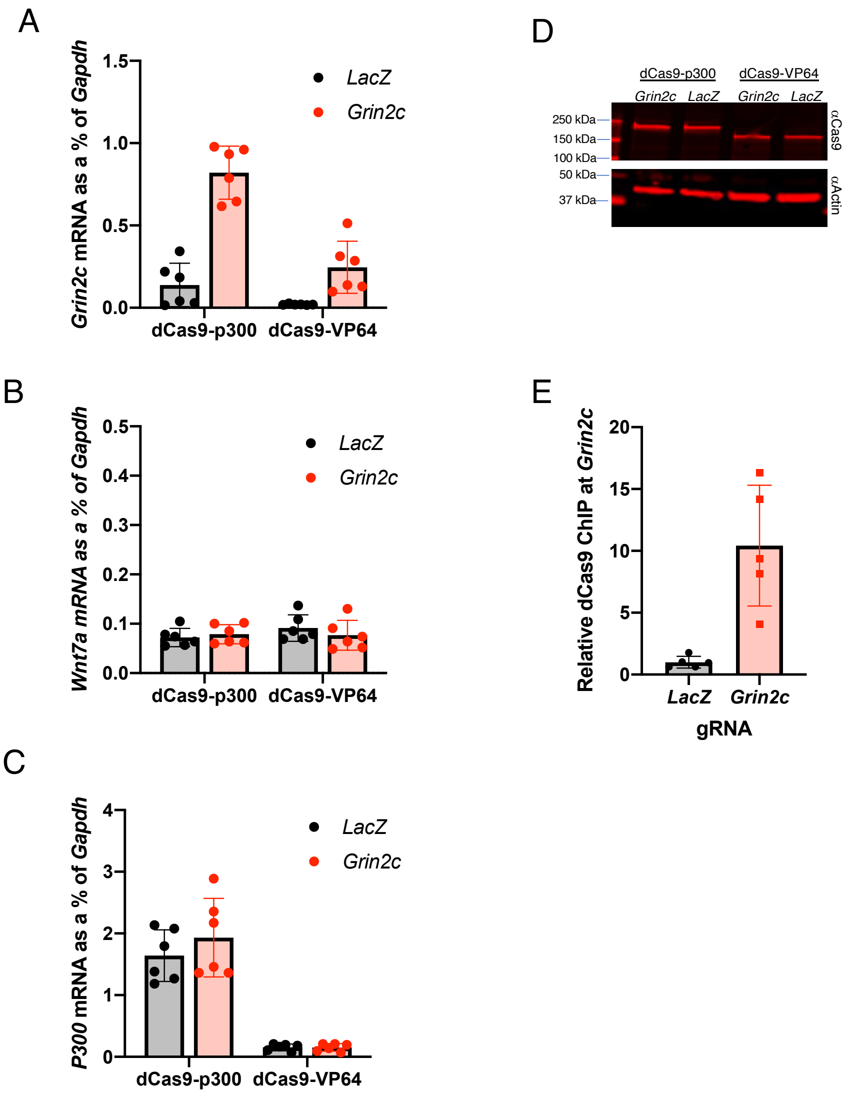


**Supplemental Figure 7. Comparison of dCas9$^{VP64}$ and dCas9$^{p300}$ activation of *Grin2c* expression in cerebellar granule neurons.** A) qRT-PCR comparing *Grin2c* mRNA in CGNs transduced with lentivirus encoding either dCas9$^{VP64}$ or Cre to induce the dCas9$^{p300}$ transgene, as well as control *LacZ* or *Grin2c* promoter-targeting gRNAs. Two-way ANOVA was significant for gRNA $F(1,20)=72.26$ $p<0.0001$, dCas9 $F(1,20)=42.08$ $p<0.001$ and gRNA x dCas9 interaction $F(1,20)=18.21$ $p=0.0004$. n=6/condition. *Grin2c* vs *LacZ* gRNA for dCas9$^{p300}$ $p<0.0001$ and for dCas9$^{VP64}$ $p=0.43$. dCas9$^{p300}$ vs dCas9$^{VP64}$ for *Grin2c* $p<0.0001$. B) qRT-PCR for *Wnt7a* mRNA in the same samples as (A) as a measure of maturity of the neurons. Two-way ANOVA showed no significant effect of gRNA $F(1,20)=0.16$ $p=0.69$ of dCas9 $F(1,20)=0.74$ $p=0.40$. C) qRT-PCR for human *P300* mRNA in the same samples from (A) to demonstrate transgene induction by *Cre* addition. Two-way ANOVA was significant for dCas9 $F(1,20)=108.9$ $p<0.0001$ but not gRNA $F(1,20)=0.86$ $p=0.36$. D) Representative western blot of dCas9 expression in cerebellar granule neurons transduced as in (A). Actin is shown as a loading control. Molecular weight (MW) markers shown on left. E) ChIP-qPCR for dCas9 from CGNs of the *dCas9$^{p300}$* transgenic mice transduced with *Cre* and the *LacZ* or *Grin2c*-gRNAs. *Grin2c* signal in the pulldown was normalized to *Gapdh* in the same sample as a control for sample processing. *Grin2c* vs *LacZ* $p=0.0026$, n=5/condition.

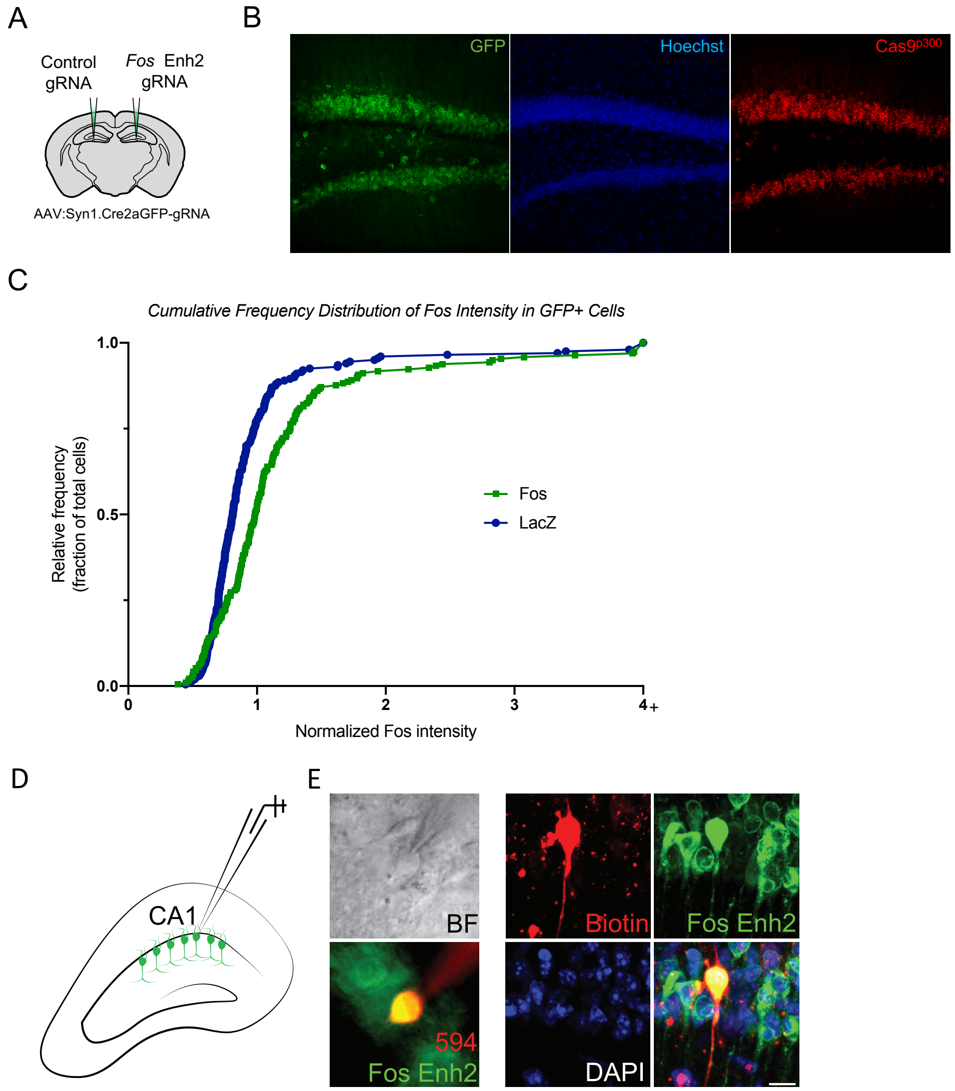


**Supplemental Figure 8: Validation of *in vivo* stereotaxic viral injection into the brain to activate *dCas9*$^{p300}$ expression and neuronal gene regulation.** A) Contralateral injection strategy to compare two treatments in the same animal. B) Delivery of the AAV vector encoding Cre and *GFP* under the control of the *Syn1* promoter and a gRNA expression cassette by injection into the hippocampus of *Rosa26:LSL-dCas9*$^{p300}$ mice induces d*Cas9* expression in most neurons. Scale bar = 50µm. C) Cumulative frequency distribution of FOS fluorescence intensity for all GFP+ cells, normalized to the average FOS intensity in the paired control. *LacZ*-gRNA: n=191 cells, *Fos* Enh2-gRNA: n=192 cells from 3 animals. $p<0.001$ by K-S test. D) Schematic of current clamp recording of CA1 neurons. E) Images confirming CA1 neurons that were recorded are positive for both biotin and GFP+, with GFP being an indicator that AAV has transduced the recorded cell and biotin introduced through the patch pipette after recording. BF, bright field, scale bar = 10µm.

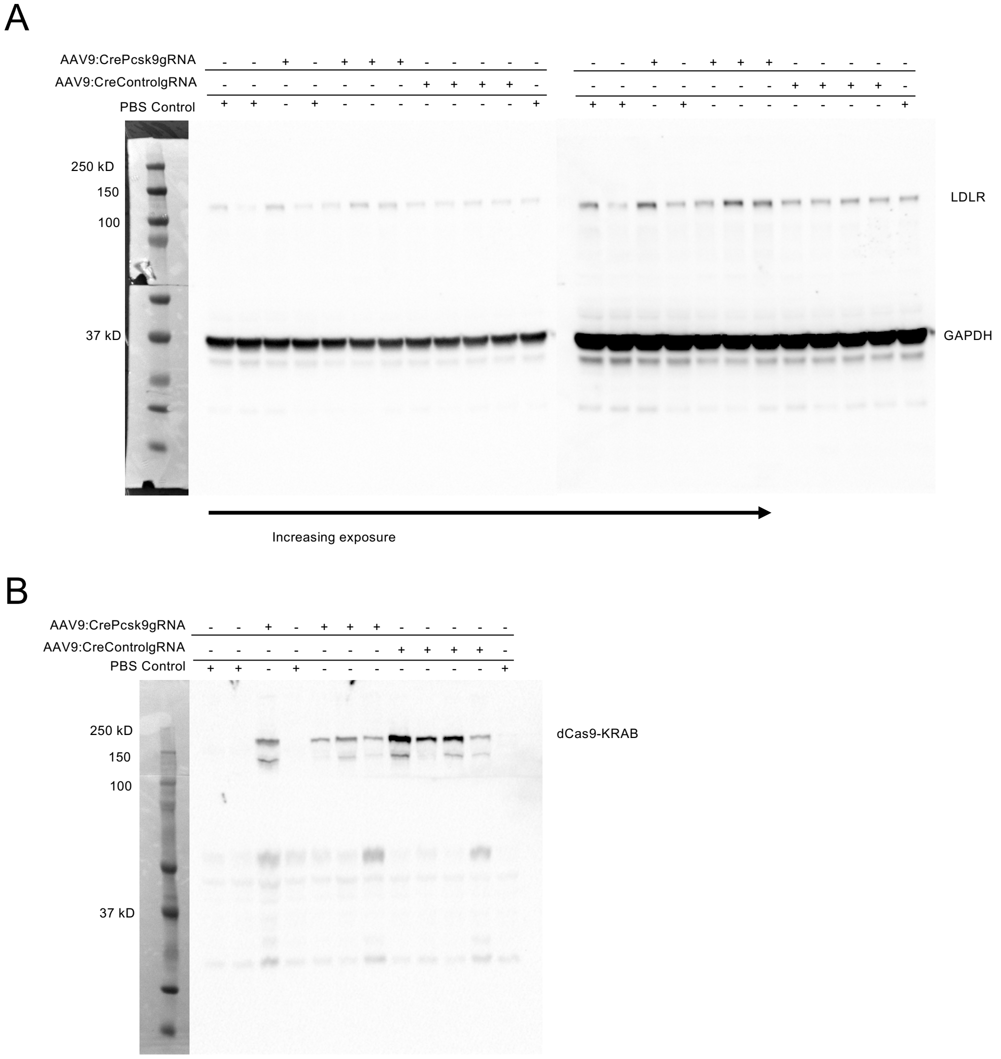


**Supplemental Figure 9: Western blots showing *dCas9*$^{KRAB}$ expression and increase in LDLR upon repression of *Pcsk9*.** A) Western blots visualized with low (left) and high (right) exposure times showing increased LDLR protein in the 3 samples with *dCas9*$^{KRAB}$ targeted to the promoter of *Pcsk9*, but not in control-gRNA or saline treated controls (n=4 per group). B) Western blots from the same mice showing *dCas9*$^{KRAB}$ expression in animals that received AA9:C Bh.Cre-gRNA (n=4 per group).

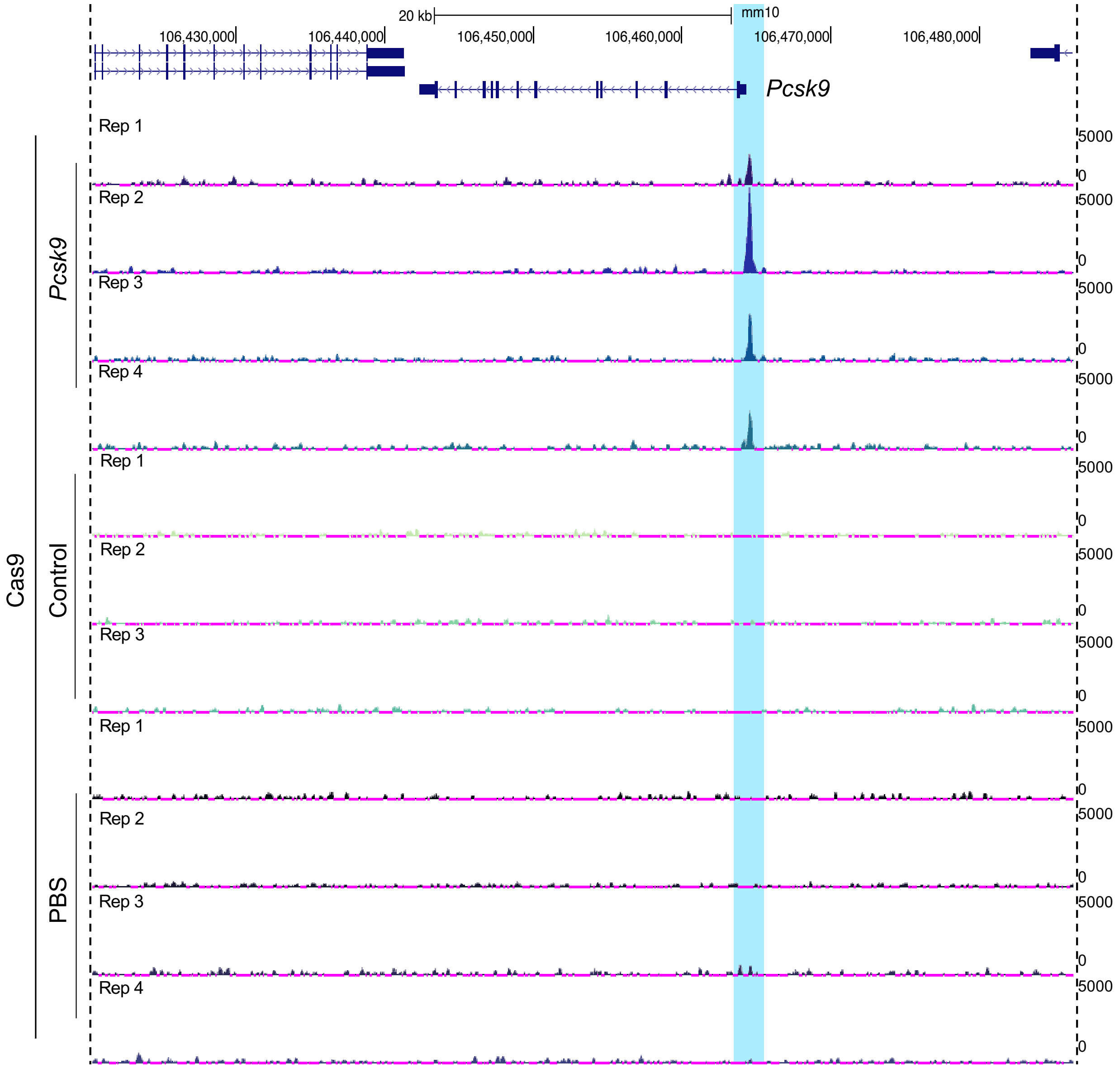


**Supplemental Figure 10: Specificity of *dCas9^KRAB^* binding the *Pcsk9* promoter *in vivo* in the liver of *Rosa26:LSL-dCas9^KRAB^* mice following systemic administration of AAV9 encoding *Cre* and a gRNA.** Genome browser tracks showing FLAG ChIP-seq signal in the liver of *dCas9^KRAB^* mice 14 days post-treatment with AAV9:Cbh.Cre-*Pcsk9*.gRNA, AAV9:Cbh.Cre-*control*.gRNA, or PBS. Highlighted regions show the area where *dCas9^KRAB^* is targeted to the promoter of *Pcsk9*. Specific signal is found only in the *Pcsk9*-gRNA treated animals.

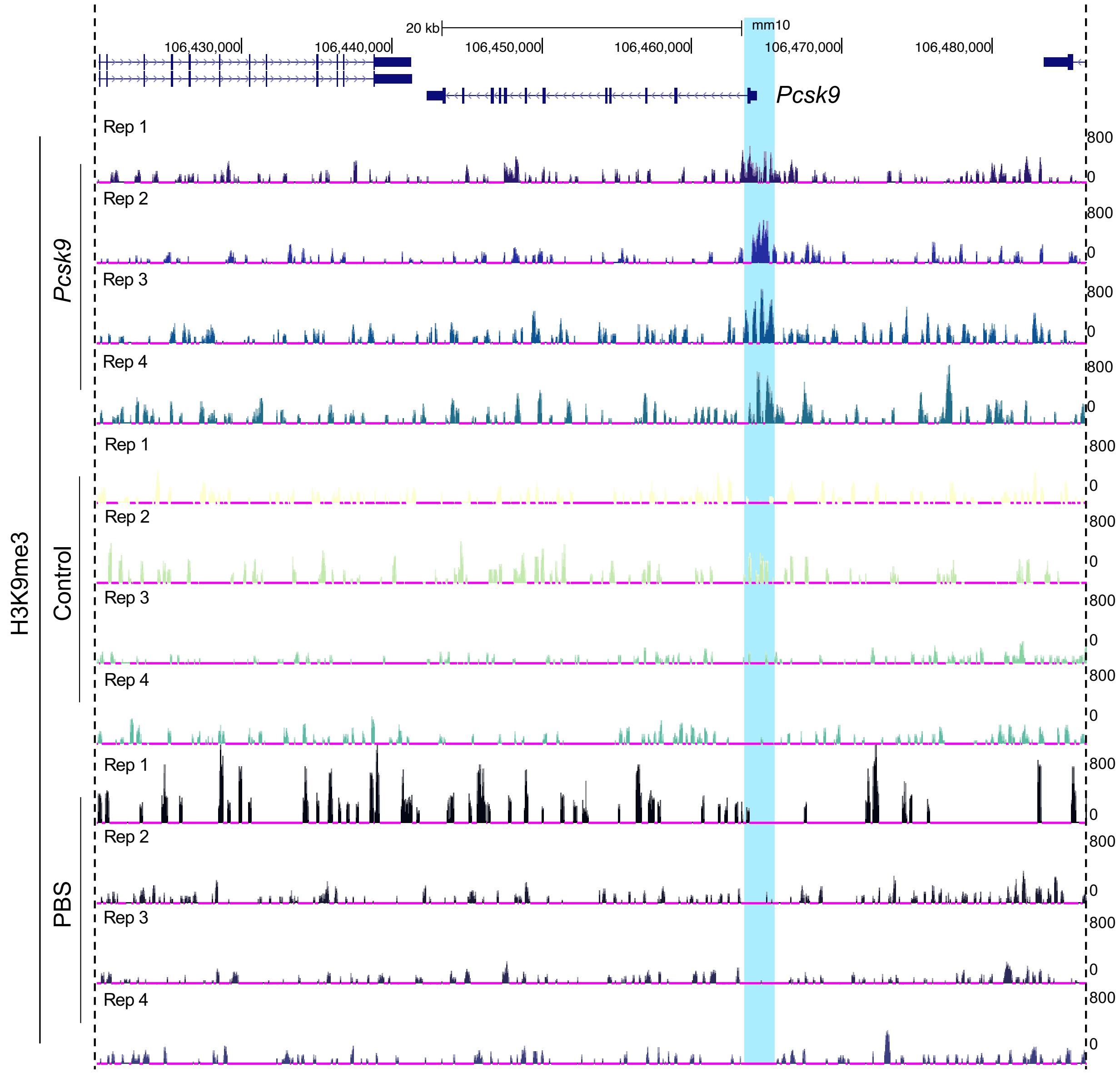


**Supplemental Figure 11: H3K9me3 enrichment at the *Pcsk9* promoter *in vivo* in the liver of Rosa26:LSL-*dCas9*$^{KRAB}$ mice following systemic administration of AAV9 encoding *Cre* and a gRNA.** Genome browser tracks showing H3K9me3 ChIP-seq signal for from livers of *dCas9*$^{KRAB}$ mice 14 days post-treatment with AAV9:Cbh.Cre-*Pcsk9.*gRNA, AAV9:CBh.Cre-*control.*gRNA, or PBS. Highlighted regions show the area where dCas9$^{KRAB}$ is targeted to the promoter of *Pcsk9*. Enrichment of H3K9me3 is noticeable in around the dCas9$^{KRAB}$ binding site in the *Pcsk9*-gRNA treated animals.

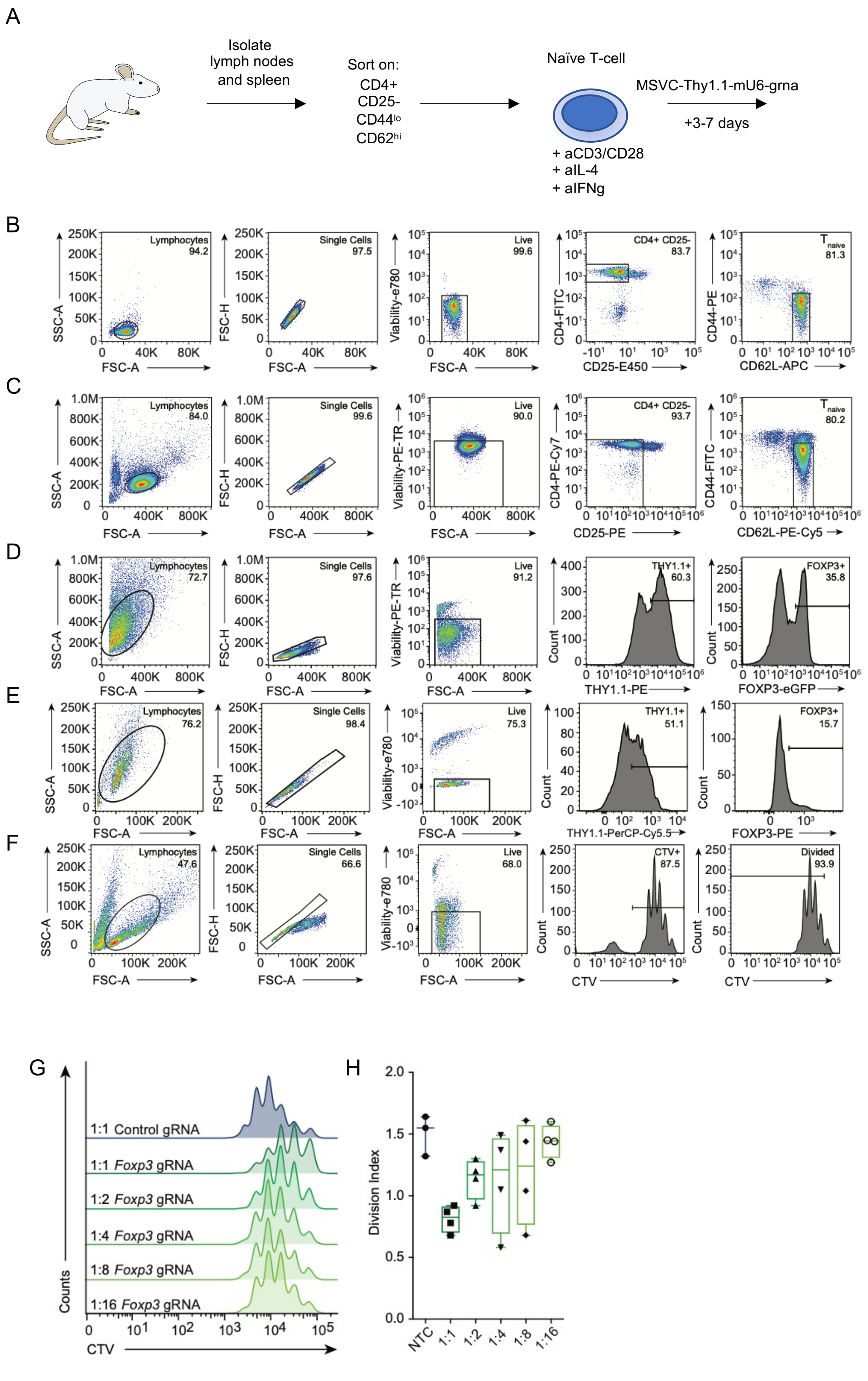


**Supplemental Figure 12: Manipulation of gene expression in T cells from *dCas9*$^{p300}$ and *dCas9*$^{KRAB}$ mice.** A) Schematic of T cell isolation strategy for the *Rosa26:LSL-dCas9*$^{p300}$ or *dCas9$^{KRAB}$* mice. B) Representative sort strategy for the purification of naïve CD4+ Tcells using the Beckman Culture Astiros or B) SONY SH800S sorter (Lymphocytes / Single Cells / Live cells / CD4+ / CD25- / CD62L$^{hi}$ / CD44$^{lo}$). **C)** Representative sort strategy for dCas9$^{p300}$-induced Tregs used in all experiments (Lymphocytes / Single Cells / Live cells / THY1.1+ / FOXP3-eGFP+). **D-E)** Gating strategy used for analyzing all *Foxp3* activation and repression experiments and to interpret the effect of targeting *Foxp3* with D) dCas9$^{p300}$ in Th0 cells or E) dCas9$^{KRAB}$ in iTreg cells (Lymphocytes / Single Cells / Live cells / THY1.1+ / FOXP3+). **F)** Gating strategy used for analysis of all suppression assays in order to determine the proliferation of Tconv when in co-culture with dCas9$^{p300}$-induced Tregs (Lymphocytes / Single Cells / Live cells / Cell Trace Violet+ / Divided). **G)** Cell proliferation traces from *in vitro* assays of suppression of T cell proliferation when co-cultured with T cells expressing control non-targeting gRNA or various dilutions of T cells expressing the *Foxp3*-targeting gRNA. **H)** Quantification of division index from this suppression assay, showing a significant difference between cells treated with the control-targeting gRNA and *Foxp3*-targeting gRNA (*$p<0.0346$, ANOVA with Dunnett's post-hoc, N=3 or 4).

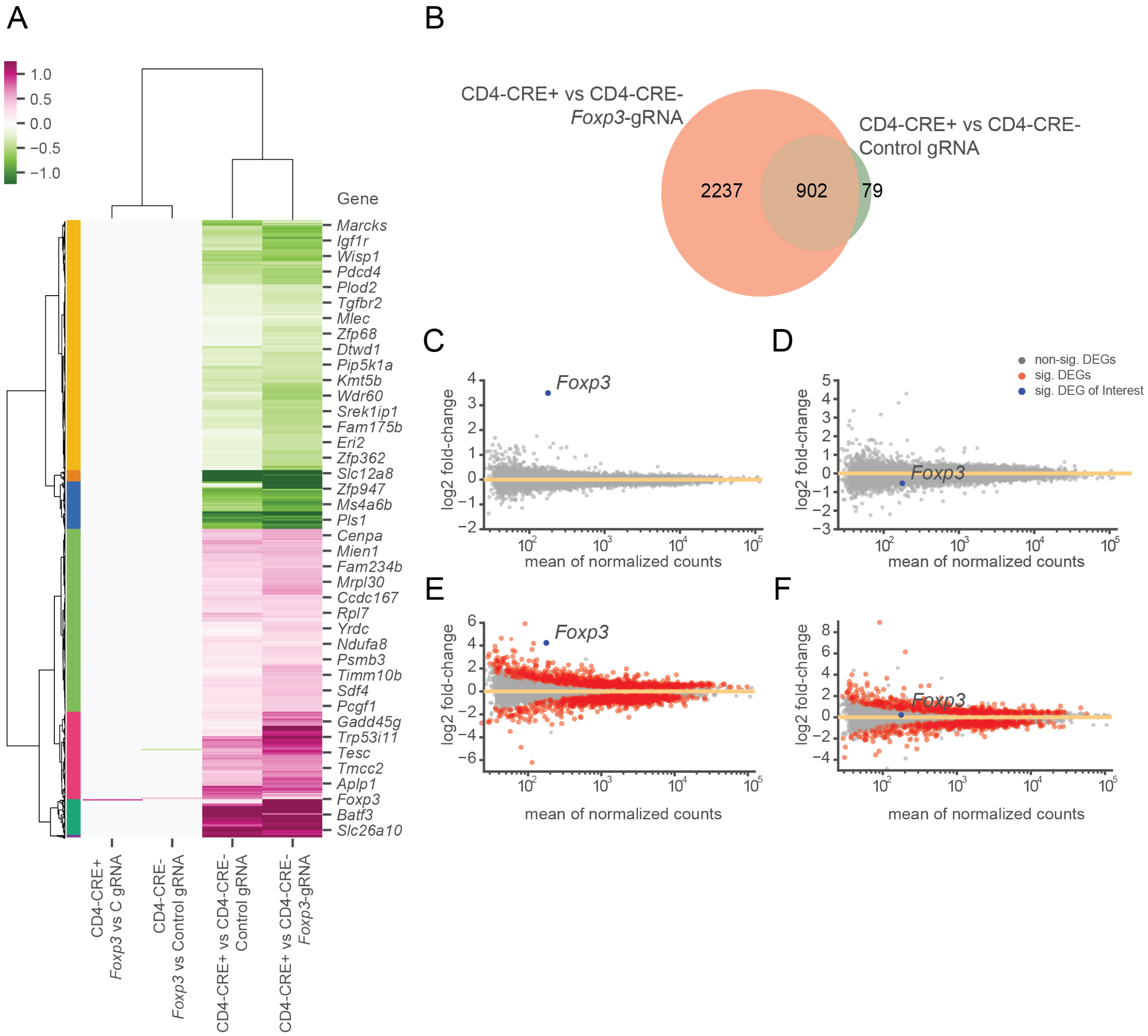


**Supplemental Figure 13: Genome wide-specificity of gene regulation by *dCas9*$^{p300}$ determined by RNA-seq in Th0 cells.** A) Heatmap visualization and hierarchical clustering of $\log_2$(fold-change) in gene expression comparing Th0 cells from *Rosa26:LSL-dCas9*$^{p300}$ mice crossed with or without *Cd4:Cre* and treated with the indicated gRNAs. B) The intersection between genes found to be differentially expressed when comparing Th0 cells with or without *Cd4:Cre* and treated with the *Foxp3*-targeting gRNA or with the control non-targeting gRNA. C-F) MA plots depicting $\log_2$(fold-change) gene expression changes for C) *Cd4:Cre*+ *Foxp3*-gRNA vs *Cd4:Cre*+ control-gRNA; D) *Cd4:Cre*- *Foxp3*-gRNA vs *Cd4:Cre*- control-gRNA; E) *Cd4:Cre*+ *Foxp3*-gRNA vs *Cd4:Cre*- *Foxp3*-gRNA; F) *Cd4:Cre*+ control-gRNA vs *Cd4:Cre*- control-gRNA. Significant differentially expressed genes are in red, FDR < 0.01; *Foxp3* target gene is labeled in blue.
